# A sensitive red/far-red photoswitch for controllable gene therapy in mouse models of metabolic diseases

**DOI:** 10.1101/2024.09.30.615971

**Authors:** Longliang Qiao, Lingxue Niu, Meiyan Wang, Zhihao Wang, Deqiang Kong, Guiling Yu, Haifeng Ye

**Affiliations:** Shanghai Frontiers Science Center of Genome Editing and Cell Therapy, Biomedical Synthetic Biology Research Center, Shanghai Key Laboratory of Regulatory Biology, Institute of Biomedical Sciences and School of Life Sciences, East China Normal University, Dongchuan Road 500, Shanghai 200241, China; School of Medicine, Shanghai University, Shanghai 200444, China; Chongqing Key Laboratory of Precision Optics, Chongqing Institute of East China Normal University, Chongqing 401120, China

## Abstract

Red light optogenetic systems are in high demand for the precise control of gene expression for gene- and cell-based therapies. Here, we report a <u>red</u>/far-red light-inducible <u>p</u>hotoswitch (REDLIP) system based on the chimeric photosensory protein FnBphP (Fn-REDLIP) or PnBphP (Pn-REDLIP) and their interaction partner LDB3, which enables efficient dynamic regulation of gene expression with a timescale of seconds without exogenous administration of a chromophore in mammals. We used the REDLIP system to establish the REDLIP-mediated CRISPR-dCas9 (REDLIP_cas_) system, enabling optogenetic activation of endogenous target genes in mammalian cells and mice. The REDLIP system is small enough to support packaging into adeno-associated viruses (AAVs), facilitating its therapeutic application. Demonstrating its capacity to treat metabolic diseases, we show that an AAV-delivered Fn-REDLIP system achieved optogenetic control of insulin expression to effectively lower blood glucose levels in type 1 diabetes model mice and control an anti-obesity therapeutic protein (thymic stromal lymphopoietin, TSLP) to reduce body weight in obesity model mice. REDLIP is a compact and sensitive optogenetic tool for reversible and non-invasive control that can facilitate basic biological and biomedical research.

## INTRODUCTION

Given the increasing demand for precision medicine, developing technologies that can precisely coordinate the release kinetics of therapeutic agents has been an area of active research^1–6^. Many precise control systems have been designed to sense and respond to chemical cues^7–11^ and physical cues, including light^12–17^, magnetic and mechanical forces^18^, temperature^19^, and electricity^20, 21^. Light has features including high spatiotemporal resolution, excellent adjustability and reversibility, and optogenetics, and has been widely used for the light-mediated manipulation of biological processes in living organisms, which has paved the way for precision interventions in gene- and cell-based therapies^22, 23^.

For many optogenetic applications, red/far-red light is preferable to other wavelengths^24, 25^, due to its advantages, including deep tissue penetration, low light scattering, and low photo-cytotoxicity^23^. Progress has recently been made toward developing red/far-red light-responsive optogenetic tools for the precise, traceless, and remote control of therapeutic gene expression in many experimental disease models^13, 14, 24–29^. Various red-shifted photoresponsive proteins have been characterized and engineered to provide flexible options for optogenetic applications based on red light (RL) illumination. The red light-responsive systems based on plant phytochrome B and phytochrome interacting factor 3 (PIF3) or 6 (PIF6) from *Arabidopsis thaliana* have been applied for transcriptional control^30, 31^, cell signaling^32, 33^, and protein localization^34^. However, these systems are limited by the requirement of supplying an exogenous chromophore and by their low packaging capacity for use with adeno-associated viral (AAV) vectors^23^. Our recently reported red/far-red light-mediated and minimized ΔPhyA-based photoswitch (REDMAP) system showed fast ON/OFF kinetics, and a relatively small construct size suitable for AAV virus packaging^24^. However, its applicability, especially for long-term controllable gene therapy, is still limited by the need for phycocyanobilin (PCB), which must be provided through exogenous administration or engineering cells to introduce PCB biosynthesis enzymes^24^.

Another class of bacteriophytochrome-based optogenetic systems that do not require exogenous supplementation of biliverdin has been developed for optogenetic manipulation in mammalian cells and mice^14, 35–37^. Our previously reported bacterial phytochrome BphS optical controllable system exhibited strong transcriptional activation but required continuous 4-6 hours of far-red light (FRL) illumination^14, 28^. Moreover, its multiple circuit components exceed the AAV packaging size. Recently, BphP1/PpsR2 (engineered QPAS1) derived from *Rhodopseudomonas palustris*^35^ and IsPadC-PCM from *Idiomarina sp. A28L*^36^ have been developed as near-infrared (NIR) optogenetic systems to precisely control gene expression in a timescale of minutes. However, they are limited by low transcriptional (about 6-fold) activation in deep tissues of mice^35, 36^. Recently, MagRed system based on *Dr*BphP derived from *Deinococcus radiodurans* and its photo-state-specific binder (named Aff6_V18FΔN) that uses split-protein reassembly to control gene expression with red light illumination^38^. However, this system has low transcriptional activation in mammalian cells and requires repetitive pulses of illumination over 24 hours^38^, indicating the low possibility for controllable gene- and cell-based therapy *in vivo*. A recently reported BICYCL-Green system employs a cyanobacteriochrome GAF domain from *Acaryochloris marina* (*AM1_C0023g2*)^37^. This system relies on green light to terminate transgene expression, potentially posing a drawback for *in vivo* applications^37^.

In this study, we sought to develop a <u>red</u>/far-red light-inducible <u>p</u>hotoswitch (REDLIP) system based on the chimeric photosensory proteins PnBphP (Pn-REDLIP) or FnBphP (Fn-REDLIP) and their common interaction partner LDB3. We first engineered PnBphP or FnBphP by fusing the N-terminal extension (NTE) of phytochrome A (PhyA)^39^ or the fungal phytochrome FphA derived from *Aspergillus nidulans*^40^ to the N-*terminus of Dr*BphP-PCM, where NTE was reported to exert a stabilizing effect on the active far-red light-absorbing state (Pfr)^40, 41^. Under red light (RL, 660 nm) illumination, PnBphP or FnBphP interacts with their common interaction partner LDB3 to allow for the precise control of gene expression; when illuminated with far-red light (FRL, 780 nm), FnBphP or PnBphP dissociate from LDB3 to terminate transgene expression^42^. These genetically encoded REDLIP systems require no external biliverdin, exhibit fast ON/OFF kinetics (10 seconds RL illumination and 1-minute FRL illumination), strong activation of target gene expression (FnBphP: 65-fold; PnBphP: 106-fold), and a dose-dependent response to red/far-red light. We also demonstrate that the REDLIP-mediated CRISPR-dCas9 (REDLIP_cas_) system enabled high induction (FnBphP: 328-fold; PnBphP: 1158-fold) of multiple endogenous genes under 10 seconds RL illumination. Moreover, the Fn-REDLIP components can be delivered to mouse muscles by AAV vectors and are used to induce RL-dependent luciferase reporter expression in mice. We also demonstrate that an AAV-delivered Fn-REDLIP system could dynamically regulate insulin expression to lower blood glucose in streptozotocin (STZ)-induced type 1 diabetic model mice and to control thymic stromal lymphopoietin (TSLP) expression^43^ to treat obesity in high-fat diet (HFD)-fed model mice. Our study thus illustrates a powerful tool for the optogenetic control of transgene expression and user-defined endogenous gene activation, enabling reversible and non-invasive control that can facilitate basic biological and biomedical research.

## RESULTS

### Design, construction, and optimization of the REDLIP system

To generate a <u>red</u>/far-red light-inducible <u>p</u>hotoswitch (REDLIP) system, we focused on the bacterial phytochrome photoreceptor *Dr*BphP from *Deinococcus radiodurans*. *Dr*BphP uses biliverdin to respond to light signals, and since this molecule is ubiquitous and endogenous in mammalian cells and animals, there is no need to administer an exogenous chromophore^35, 38, 42^. Based on a previous study reporting that PhyA’s NTE influences photoresponse properties by stabilizing the active Pfr state ^39, 40, 44^, we initially engineered chimeric photosensory proteins BphP (PnBphP or FnBphP) by fusing the N-terminal extension (NTE) of phytochrome A (PhyA) from *Arabidopsis thaliana* or the fungal phytochrome FphA from *Aspergillus nidulans* to the N-*terminus of Dr*BphP-PCM.

Next, we constructed three types of REDLIP systems (Pn-REDLIP, Fn-REDLIP, Dr-REDLIP), in which the PnBphP, FnBphP, or the original *Dr*BphP was fused to Gal4, a yeast DNA binding domain, to form a hybrid DNA binding protein (Gal4-PnBphP, Gal4-FnBphP, or Gal4-*Dr*BphP). Their common interacting partner LDB3 was fused to a hybrid *trans-*activator p65-HSF1 [the 65-kDa *trans-*activator subunit of NF-κB (p65) and heat shock factor 1 (HSF1) transactivation domain] to form the light-inducible *trans-*activator LDB3-p65-HSF1. Upon RL illumination, PnBphP, FnBphP, or *Dr*BphP interacts with LDB3, enabling it to bind to an RL-specific chimeric promoter (P_RL_, 5×UAS-P_TATA_) via the Gal4 domain to initiate transgene expression. Upon exposure to FRL, the BphPs dissociate from LDB3, thereby terminating target transgene expression (**Fig 1a, b**). In addition, we evaluated the impact of the different BphPs fused to Gal4 and found the Fn-REDLIP system efficiently activated SEAP reporter expression upon RL illumination, and the Pn-REDLIP system significantly decreased leakiness in the dark compared to the Dr*-*REDLIP system (**Supplementary Fig. 1**).

**Fig. 1.**
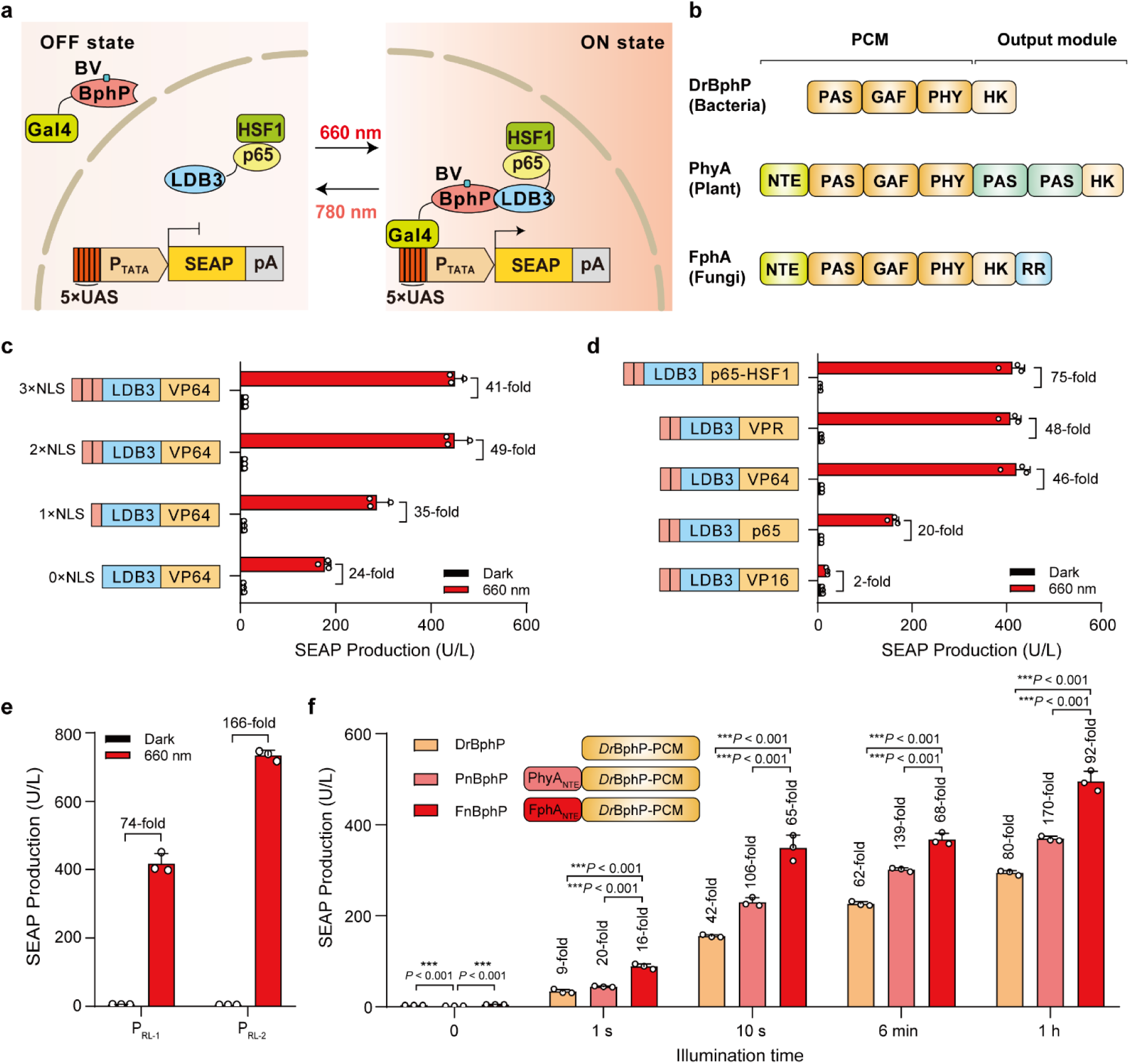
Construction and optimization of the REDLIP system. **a,** Schematic showing the design of the REDLIP system for controlling gene expression. The BphP-PCM (photosensory core module) interaction domain nanobody LDB3 is fused to the transcriptional activation domain (p65-HSF1) to create a red light (RL, 660 nm) -dependent *trans-*activator (LDB3-p65-HSF1) driven by the human cytomegalovirus promoter (P_hCMV_). The DNA binding domain Gal4 is fused to the chimeric PnBphP (<u>P</u>hyA<u>_N_</u>_TE_-*Dr*BphP), the FnBphP (<u>F</u>phA<u>_N_</u>_TE_-*Dr*BphP), or the wild-type *Dr*BphP to create a fusion light sensor domain (Gal4-BphP) driven by P_hCMV_. When exposed to RL (660 nm), the *trans-*activator (LDB3-p65-HSF1) can specifically bind to the light sensor domain (Gal4-BphP), after which the bound complex preferentially interacts with the chimeric promoter (P_RL_, 5×UAS-P_TATA_) and initiates transgene expression. Upon exposure to FRL (780 nm), the *trans-*activator dissociates from the light sensor domain (Gal4-BphP), leading to disengagement from the promoter and terminating transgene expression. **b**, The domain organization of three phytochrome-related proteins: *Dr*BphP, PhyA, and FphA. The PCM comprises the N-terminal PAS (period/Arnt/Sim), GAF (cGMP phosphodiesterase/adenylyl cyclase/Fhl1), and PHY (phytochrome-specific) domains. At the same time, the C-terminal output module consists of a variable domain (HK, PAS, RR). HK, histidine kinase; RR, response regulator. **c,** Comparison of the performance of the copy number for the nuclear localization signal (NLS) of the RL-dependent *trans-*activator (LDB3-p65-HSF1). HEK-293T cells (6×10^4^) were co-transfected with the Gal4-FnBphP vector (pQL326), the SEAP reporter expression vector (pDL6, P_5×UAS_-SEAP-pA), and the light-inducible *trans-*activator with different copy numbers of NLS: 1NLS-LDB3-VP64 expression vector (pQL243), 2NLS-LDB3-VP64 expression vector (pQL242), 3NLS-LDB3-VP64 expression vector (pQL232), or LDB3-VP64 expression vector (pQL207), and then illuminated with RL (660 nm, 2.0 mW/cm^2^) for 24 hours; SEAP production in the culture supernatant was quantified 24 hours after illumination. **d,** Screening different transcriptional activators fused to LDB3. HEK-293T cells (6×10^4^) were co-transfected with pQL326, pDL6, and either LDB3-VP64 expression vector (pQL242), LDB3-VP16 expression vector (pQL251), LDB3-p65 expression vector (pQL250), LDB3-VPR expression vector (pQL252), or LDB3-p65-HSF1 expression vector (pNX12), and subsequently illuminated as described in c. SEAP production was quantified 24 hours after illumination. **e,** Optimization of RL-responsive P_RL-X_-driven SEAP expression. HEK-293T cells (6×10^4^) were transfected with pQL326, pNX12, and a P_RL-X_-driven SEAP reporter [pDL6 (P_RL-1_, 5×UAS-P_hCMVmin_) or pYZ430 (P_RL-2_, 5×UAS-P_TATA_)], and subsequently illuminated as described in c. SEAP production was quantified 24 hours after illumination. **f,** Comparison of the performance of different chimeric BphP constructs fused to Gal4 illuminated with RL (660 nm, 2.0 mW/cm^2^) for 1 second, 10 seconds, 6 minutes, and 1 hour. HEK-293T cells (6×10^4^) were co-transfected with pYZ430, pNX12, and either Gal4-DrBphP expression vector (pQL217), Gal4-PnBphP expression vector (pQL325), or pQL326, and then illuminated with RL (660 nm, 2.0 mW/cm^2^) for the indicated times (0-1 hour). SEAP production was quantified 24 hours after illumination. Fold change is compared to the 0-hour time point. Data in c-f are presented as means ±SD; *n* = 3 independent experiments. *P* values in f were calculated by one-way ANOVA with multiple comparisons. Detailed descriptions of the genetic constructs and transfection mixtures are provided in Supplementary Tables 1 and 4.

To optimize the induction profiles of the Fn-REDLIP system, we fused the VP64 activator [(a tetrameric repeat of the minimal *Herpes simplex-*derived *trans-*activator VP16 (herpes simplex viral protein 16)] with 1 or 2 or 3 copies of nuclear localization signal (NLS); upon RL illumination, VP64 with 2 copies of NLS (2 NLS) enhanced the fold induction (49-fold induction) compared to the dark control sample (**Fig. 1c)**. We also examined the effects of LDB3 fused with five different *trans-*activators, including VP16, p65, VP64, VP64-p65-Rta (VPR), and p65-HSF1: LDB3 fused with p65-HSF1 produced the highest induction efficiency (75-fold) of SEAP expression upon RL illumination (**Fig. 1d)**. For further improvement, we tested two RL specific chimeric promoters (P_RL-1,_ 5×UAS-P_hCMVmin;_ P_RL-2,_ 5×UAS-P_TATA_) driving SEAP expression and found that SEAP expression under the control of the P_RL2_ (5×UAS-P_TATA_) promoter resulted in a high induction (166-fold induction) (**Fig. 1e)**. We also observed that 10 seconds of RL at 2.0 mW/cm^2^ induced gene expression, achieving ∼60% of the efficiency obtained from 1 hour of illumination (**Fig. 1f)**. Compared to the Dr-REDLIP system, the Fn-REDLIP system showed the highest gene expression under RL illumination, while the Pn-REDLIP system exhibited the lowest extent of leaky activity in the dark and the highest induction efficiency (**Fig. 1f**). As such, the Pn-REDLIP system could be preferable for applications with stringent background requirements.

### Characterization of the Fn-REDLIP and Pn-REDLIP systems

We assessed the performance of the Fn-REDLIP and Pn-REDLIP systems in HEK-293T cells. First, we evaluated the kinetics of the two REDLIP systems and found that the SEAP reporter gene expression depended on the time and intensity of RL illumination (**Fig. 2a, b, and Supplementary Fig. 2a, b**) without a noticeable background increase over time under dark conditions; there was no increase in the SEAP signal within 24 hours of stopping RL illumination (**Fig. 2c and Supplementary Fig. 2c**). Remarkably, two REDLIP systems were more sensitive to red light: RL illumination for only 10 seconds at 2 mW/cm^2^ resulted in a substantial increase in SEAP expression, achieving ∼60% of the maximum by 24 h of RL illumination. SEAP signals from the two REDLIP systems were observed under RL illumination, while only minimal SEAP expression was observed under ambient light; note that no induced gene expression was observed upon exposure to ultraviolet light, blue light, green light, or far-red light (**Fig. 2d and Supplementary Fig. 2d**). The results support the Fn-REDLIP and Pn-REDLIP systems can be interpreted as specificity to one particular wavelength.

**Fig. 2.**
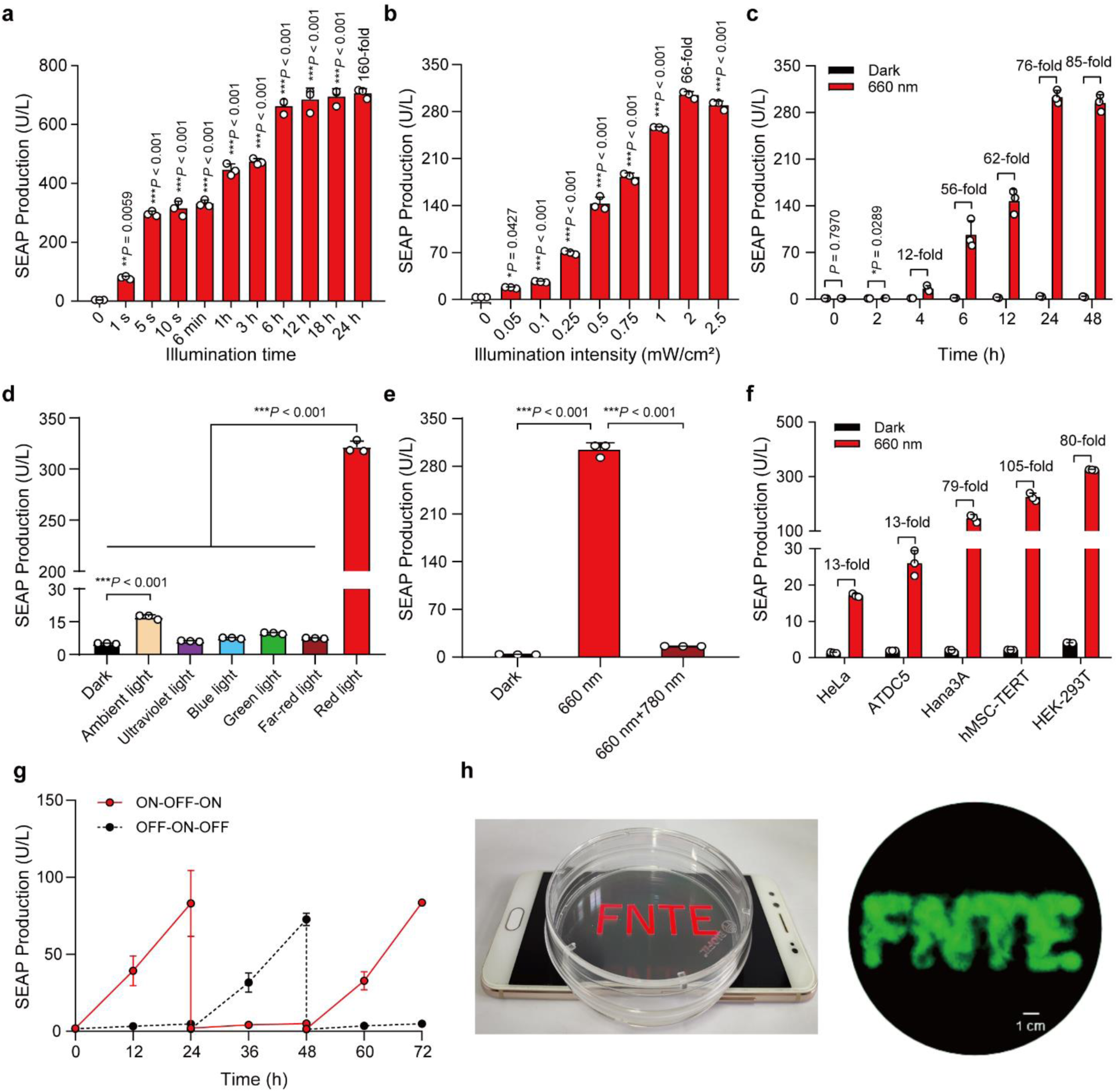
Characterization of the Fn-REDLIP system performance. **a,** Assessment of illumination time-dependent Fn-REDLIP-mediated transgene expression kinetics. HEK-293T cells (6×10^4^) were co-transfected with pQL326, pNX12, and pYZ430 at a 2:2:1 (w/w/w) ratio, and illuminated with RL (660 nm, 2.0 mW/cm^2^) for the indicated time (0-24 hours). SEAP production was quantified 24 hours after initial illumination. **b,** Illumination intensity-dependent Fn-REDLIP-mediated transgene expression kinetics. HEK-293T cells (6×10^4^) transfected as described in a were illuminated with RL (660 nm) at the indicated light intensities (0 to 2.5 mW/cm^2^) for 10 seconds. SEAP production was quantified 24 hours after illumination. **c,** Quantification of Fn-REDLIP-mediated transgene expression kinetics. HEK-293T cells (6×10^4^) transfected as described in a were illuminated with RL (660 nm, 2.0 mW/cm^2^) for 10 seconds. After illumination, SEAP production in the culture supernatant was profiled at the indicated time periods (0-48 hours). **d**, Chromatic specificity of the Fn-REDLIP system. HEK-293T cells (6×10^4^) transfected as described in (A) were illuminated with ultraviolet light (365 nm), blue light (465 nm), green light (530 nm), far-red light (780 nm), or red light (660 nm) at 2.0 mW/cm^2^ or ambient light (750 Lux) for 10 seconds. SEAP production was quantified 24 hours after illumination. **e,** The switch ON/OFF performance of the Fn-REDLIP system. Three groups of HEK-293T cells co-transfected as described in a. 24 hours after transfection, cells were illuminated with RL for 10 seconds and switched to the dark condition or 780 nm illumination at 2.0 mW/cm^2^ for 2 minutes for the “660 nm+780 nm” group. As a control, cells were kept in the dark throughout the experiment. SEAP production was quantified 24 hours after 660 nm or 780 nm illumination. **f,** Fn-REDLIP-mediated SEAP expression in the indicated mammalian cell lines. Five mammalian cell lines (6×10^4^) co-transfected as described in a were illuminated with RL (660 nm, 2.0 mW/cm^2^) for 10 seconds. SEAP production was quantified 24 hours after illumination. **g,** Reversibility of Fn-REDLIP-mediated transgene expression. HEK-293T cells (6×10^4^) transfected as described in a and were illuminated with RL (660 nm, 1.0 mW/cm^2^) for 3 seconds (ON) or kept in the dark (OFF). SEAP production was quantified every 12 hours for 72 hours; the culture medium was renewed every 24 hours. **h,** Spatial control of Fn-REDLIP-mediated transgene expression. HEK-293T cells (3×10^6^) were seeded into a 10-cm tissue culture dish and transfected with the Fn-REDLIP system and an EGFP reporter pDQ63 (P_RL_-EGFP-pA). The dish was placed onto a smartphone screen projecting a letter pattern and illuminated with RL for 15 minutes (display brightness set to 100%, 40 μW/cm^2^) (schematic, left). The fluorescence micrographs assessing EGFP production were taken 24 hours after illumination using fluorescence microscopy. Scale bar, 1 cm. Data in a-g are presented as means ±SD; *n* = 3 independent experiments. *P* values in (a, b, d, and e) were calculated by one-way ANOVA with multiple comparisons. *P* values in c were calculated using a two-tailed unpaired *t*-test. Detailed descriptions of the genetic constructs and transfection mixtures are provided in Supplementary Tables 1 and 4.

We next examined whether the Fn-REDLIP and Pn-REDLIP systems can be switched ON with RL and OFF with FRL. HEK-293T cells transfected with Fn-REDLIP or Pn-REDLIP systems were exposed to RL (660 nm, 2.0 mW/cm^2^) for 10 seconds to induce SEAP expression, followed immediately by exposure to FRL (780 nm, 1.0 mW/cm^2^) for 2 minutes. An obvious SEAP signal was only observed in cells exposed to RL illumination; no signal was evident for cells first exposed to RL and then to FRL (**Fig. 2e and Supplementary Fig. 2e**). Further, exposure to various durations and intensities of FRL (780 nm) showed that a 0.5-minute exposure time at 1.0 mW/cm^2^ was sufficient to switch OFF approximately 90% of the maximum gene expression observed in the sample exposed to RL (660 nm, 2.0 mW/cm^2^) for 10 seconds (**Supplementary Fig. 3a-e**). We also found that the two REDLIP systems were functional in five different mammalian cell types (**Fig. 2f and Supplementary Fig. 2f**). Further, the Fn-REDLIP and Pn-REDLIP systems enabled reversible control of transgene expression (**Fig. 2g and Supplementary Fig. 2g**) and exhibited spatial transgene activation upon local illumination (**Fig. 2h and Supplementary Fig. 2h**). Thus, the Fn-REDLIP and Pn-REDLIP systems represent new optogenetic tools for robust, fast, wavelength-specific, adjustable, and reversible transgene activation.

### REDLIP-controlled CRISPR-dCas9 systems for activation of endogenous gene transcription

After demonstrating that the REDLIP system induced highly efficient exogenous gene transcription under RL illumination, we investigated whether the system can control endogenous gene activation. Taking advantage of the synergistic activation mediator (SAM) system^45^, which has been confirmed to enhance dCas9-targeted endogenous gene activation by introducing sgRNA-bearing MS2 RNA aptamers, we generated an RL-induced *trans-*activator by fusing MS2 to p65-HSF1, which can be recruited by sgRNA bearing MS2 aptamers and the dCas9 complex to induce gene transcription. When FRL is turned on, the chimeric BphPs dissociate from LDB3, inactivating CRISPR-dCas9 targeted endogenous gene transcription (**Fig. 3a and Supplementary Fig. 4a**).

**Fig. 3.**
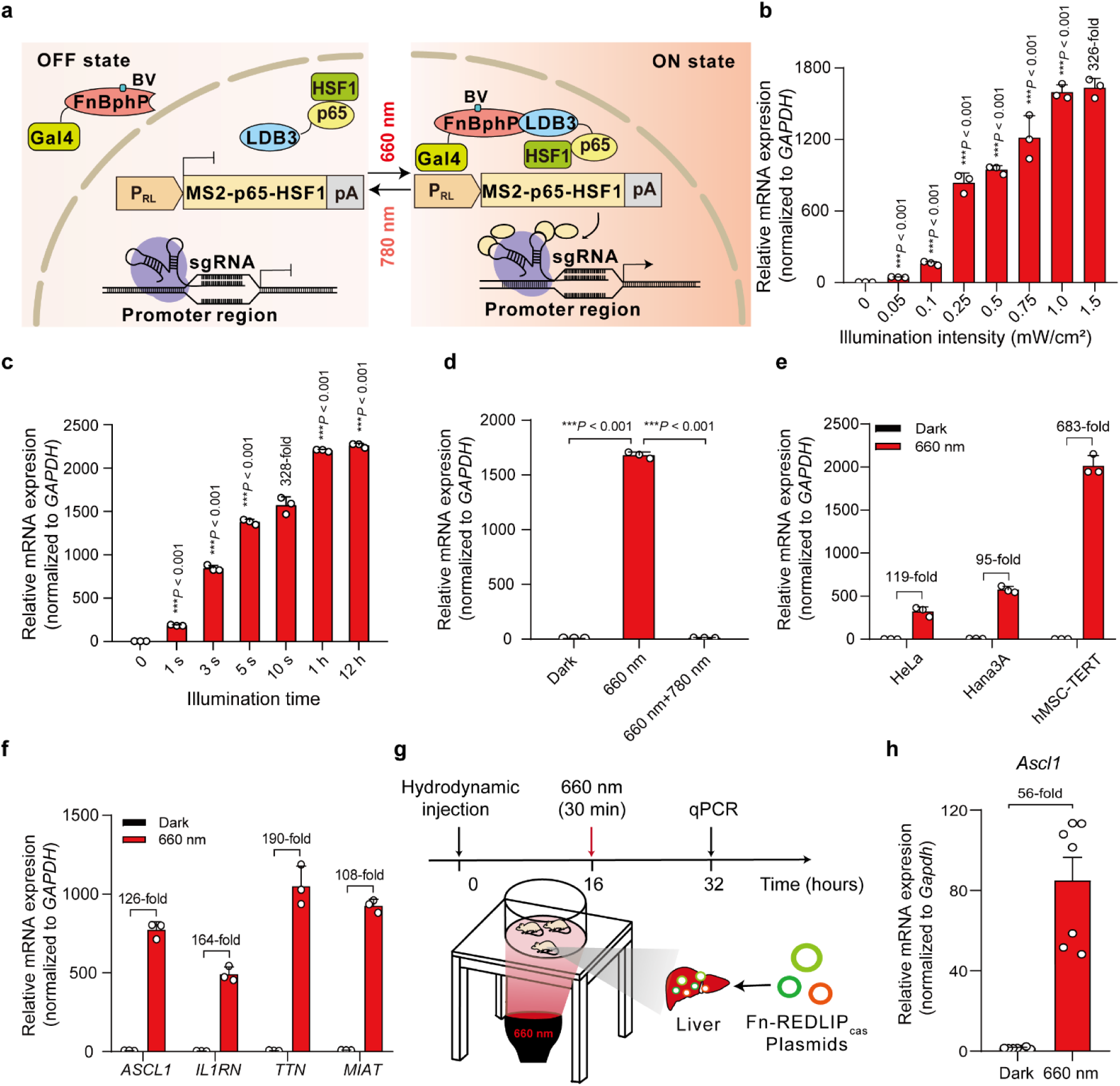
Fn-REDLIP-controlled CRISPR-dCas9 system (Fn-REDLIP_cas_) for endogenous gene activation in mammalian cells and mice. **a,** Schematic design of the Fn-REDLIP_cas_ system for transcriptional activation. The RL chimeric promoter (P_RL_) drove the expression of the *trans-* activator MS2-p65-HSF1, which can be recruited by the MS2 box of the sgRNA-dCas9 complex to activate target gene transcription under RL illumination. 780-nm light illumination can switch the Fn-REDLIP_cas_ system back to an inactive state. **b,** Illumination intensity-dependent Fn-REDLIP_cas_ system-mediated activation of endogenous gene transcription. HEK-293T (6×10^4^) cells were co-transfected with pQL326, pNX12, a P_RL_-driven MS2-p65-HSF1 expression vector (pDQ100), dCas9 expression vector (pSZ69), and two sgRNAs targeting the *RHOXF2* locus (pSZ105, P_U6_-sgRNA1*_RHOXF2_*-pA; pSZ106, P_U6_-sgRNA2*_RHOXF2_*-pA) at a 15:15:1:10:5:5 (w/w/w/w/w/w) ratio, followed by illumination with RL (660 nm) at the indicated intensities (0 to 1.5 mW/cm^2^) for 10 seconds. **c,** Illumination time-dependent Fn-REDLIP_cas_ system-mediated activation of endogenous gene transcription. HEK-293T cells (6×10^4^) were co-transfected as described in b and illuminated with RL (660 nm, 1.0 mW/cm^2^) for different time periods (0 to 12 hours). **d,** The switch ON/OFF performance of the Fn-REDLIP_cas_ system. Three groups of HEK-293T cells were transfected as described in b. 24 hours after transfection, cells were illuminated with RL for 10 seconds and switched to the dark condition or 780 nm illumination at 2.0 mW/cm^2^ for 2 minutes. As a control, cells were kept in the dark throughout the experiment. The levels of endogenous *RHOXF2* were quantified by qPCR at 24 hours after 660 nm or 780 nm illumination. *P* values in (B-D) were calculated by one-way ANOVA with multiple comparisons. **e,** The Fn-REDLIP_cas_-mediated endogenous gene activation in the indicated mammalian cell lines. Three mammalian cell lines (6×10^4^) were co-transfected as described in b and illuminated with red light (660 nm, 1.0 mW/cm^2^) for 10 seconds. **f,** The Fn-REDLIP_cas_-mediated activation of different endogenous genes. HEK-293T cells (6×10^4^) were co-transfected with Fn-REDLIP_cas_ together with two sgRNAs targeting the *ASCL1* locus, the *TTN* locus, the *IL1RN* locus, or the *MIAT* locus. Twenty-four hours after transfection, cells were exposed to red light (660 nm, 1.0 mW/cm^2^) for 10 seconds. Data in b-f are presented as relative mRNA expression levels quantified by qPCR 24 hours after illumination. All data are represented as means ±SD; *n* = 3 independent experiments. **g,** Schematic representation of the experimental procedure and time schedule for Fn-REDLIP_cas_-mediated activation of endogenous genes in mouse livers. A mixture of plasmids containing the Fn-REDLIP_cas_ system and the concatenated gRNAs and dCas9 vector pYZ561 (P_U6_-sgRNA1*_Ascl1_*-pA::P_U6_-sgRNA2*_Ascl1_*-pA::P_hCMV_-dCas9-pA) were hydrodynamically injected into mice. Sixteen hours after injection, the mice were illuminated with red light (660 nm, 20 mW/cm^2^) for 30 min, or kept in the dark. At 16 hours after illumination, mice were euthanized, and their livers were harvested for qPCR analysis of *Ascl1* expression. **h,** qPCR analysis of the *Ascl1* activation with the Fn-REDLIP_cas_ system in mice. Data in h are presented as means ±SEM (*n* = 7 mice). Detailed descriptions of the genetic constructs and transfection mixtures are provided in Supplementary Tables 1 and 4.

To assess the REDLIP-mediated CRISPR-dCas9 systems (Fn-REDLIP_cas_ and Pn-REDLIP_cas_) for activating transcription of the Rhox homeobox family member 2 (*RHOXF2*) gene, we co-transfected HEK-293T cells with the Fn-REDLIP_cas_ or the Pn-REDLIP_cas_ system and two sgRNAs targeting the *RHOXF2* locus. We found that the transcriptional activation of *RHOXF2* depended on the intensity and time of RL illumination (**Fig. 3b, c, and Supplementary Fig. 4b, c**). Note that a very short (10 seconds) illumination efficiently activated *RHOXF2* transcription using the Pn-REDLIP_cas_ system (1158-fold induction) with no apparent leaky activity in the dark or using the Fn-REDLIP_cas_ system (328-fold induction) with a higher level of gene transcription (**Supplementary Fig. 4c and Fig. 3c**). Moreover, our two REDLIP_cas_ systems could activate *RHOXF2* transcription under RL illumination and deactivate transcription under RL and then FRL illumination (**Fig. 3d and Supplementary Fig. 4d**).

Supporting the broad application potential of the two REDLIP_cas_ systems, we transfected the two REDLIP_cas_ systems into three different cell lines, respectively, and found that they can activate endogenous *RHOXF2* in the tested cells (**Fig. 3e and Supplementary Fig. 4e**). In addition, we successfully deployed them to activate the transcription of multiple endogenous genes using four pairs of sgRNAs, with each pair targeting the promoter of a single gene, including achaete-scute homolog 1 (*ASCL1)*, interleukin 1 receptor antagonist *(IL1RN*), titin (*TTN*), or myocardial infarction associated transcript (*MIAT*) (**Fig. 3f and Supplementary Fig. 4f**).

We next examined whether the REDLIP_cas_ system can activate endogenous gene transcription in mice under RL illumination. We transiently transfected the Fn-REDLIP_cas_ system and two sgRNAs targeting the *ASCL1* locus into C57BL/6 mice using hydrodynamic injection via the tail vein. Sixteen hours after injection, the abdomens of the transfected mice were illuminated with RL for 30 minutes (660 nm, 20 mW/cm^2^) (**Fig. 3g**). qPCR showed that the *ASCL1* level was significantly up-regulated (56-fold) in illuminated mouse livers compared to dark control mouse livers (**Fig. 3h**). These results demonstrate that the Fn-REDLIP_cas_ system can activate the transcription of user-defined endogenous genes in mammalian cells and mice upon illumination with a noninvasive LED.

### Optogenetic control of gene expression using AAV-delivered Fn-REDLIP system in mice

For gene transcription activation *in vivo*, we also transiently transfected wild-type C57BL/6 mice with Gal4-FnBphP or Gal4-PnBphP or Gal4-*Dr*BphP and LDB3-p65-HSF1 and luciferase reporter plasmids (5×UAS-P_TATA_-Luciferase-pA) using hydrodynamic injection via the tail vein. Sixteen hours after injection, the transfected mice were illuminated with RL (660 nm, 20 mW/cm^2^) for different time periods (0 to 60 minutes) **(Fig. 4a and Supplementary Fig. 5a).** Luciferase reporter expression was measured using an *in vivo* imaging system: mice illuminated with RL exhibited significantly higher luciferase expression compared with the dark control mice. Moreover, the *in vivo* transgene expression induced by these systems was exposure-time dependent (**Fig. 4b, c, Supplementary Fig. 5b, c and 6a-c)**. Strikingly, RL illumination for 1 minute was sufficient to induce luciferase expression in the Fn-REDLIP or Pn-REDLIP system-equipped mice. In contrast, a bioluminescence signal was observed upon RL illumination for a much longer illumination time (30 minutes) in mice bearing the Dr-REDLIP system. Moreover, the mice injected with the Fn-REDLIP system showed stronger bioluminescence signal intensities in livers compared to mice injected with the Pn-REDLIP system, and the mice injected with the Dr-REDLIP system (**Fig. 4c and Supplementary Fig. 5b, c)**. These results demonstrate that the Fn-REDLIP system can efficiently regulate transgene expression in deep organ tissues of mice, giving it potential for applications *in vivo*.

**Fig. 4.**
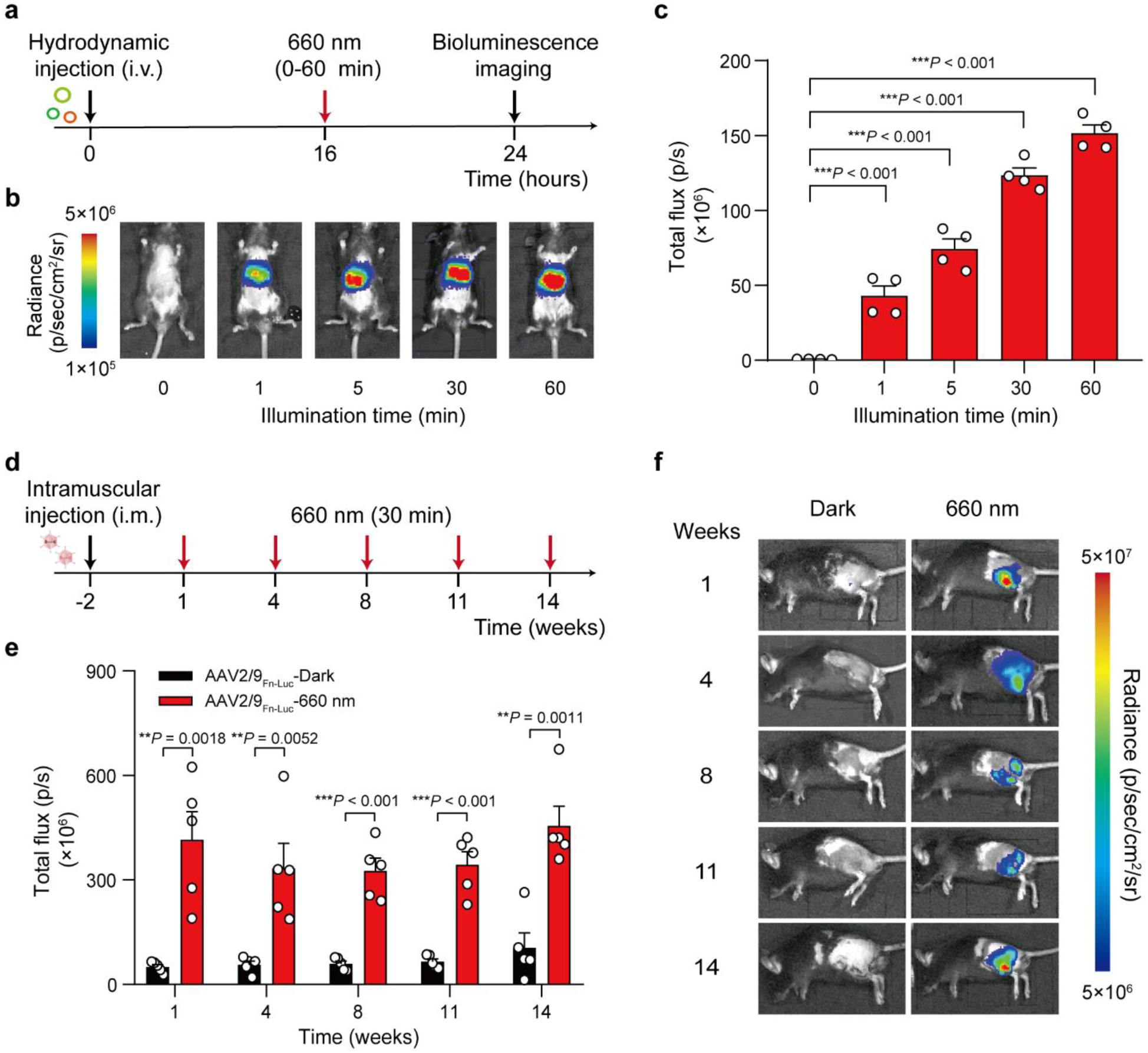
Fn-REDLIP-mediated transgene expression in mice. **a,** Schematic representation of the experimental procedure and the time schedule for Fn-REDLIP-mediated transgene expression in mouse livers. **b, c,** Exposure time-dependent Fn-REDLIP-mediated transgene expression kinetics in mice. Mice were hydrodynamically injected (tail vein) with Fn-REDLIP-encoding plasmids. At 16 hours post-injection, the mice were illuminated with RL (660 nm, 20 mW/cm^2^) for the indicated time periods (0-60 minutes). The bioluminescence signal was quantified 8 hours after illumination using an *in vivo* imaging system. **d-f,** Adeno-associated virus (AAV)-serotype of 2/9 delivery of the Fn-REDLIP-mediated transgene expression in mice. **d,** Schematic representation of the experimental procedure and time schedule for the AAV2/9 delivery of Fn-REDLIP-mediated transgene expression in the mouse left gastrocnemius muscle. Mice were intramuscularly injected with a mixture of AAVs encoding the Fn-REDLIP system [(pQL382 (ITR-P_EMS_-Gal4-FnBphP-pA-ITR) and pNX137 (ITR-P_EMS_-LDB3-p65-HSF1-pA-ITR)] and the luciferase reporter pQL271 (ITR-P_RL_-Luciferase-pA-ITR) at an AAV titer of 2×10^11^ Vector Genomes (vg). After 2 weeks, mice were illuminated at an intensity of 20 mW/cm^2^ for 30 minutes once every three or four weeks. **e,** Bioluminescence was quantified 8 hours after illumination. **f,** Bioluminescence measurements of long-term activated luciferase expression of the AAV-delivered Fn-REDLIP system in mice under red light illumination. Data in c, f are presented as means ±SEM (*n* = 4 or 5 mice). *P* values in c were calculated by one-way ANOVA with multiple comparisons. *P* values in e were calculated using a two-tailed unpaired *t*-test. Detailed bioluminescence images of the mice are provided in Supplementary Figs. 6a, 8.

AAV vectors are frequently used in biological research and for gene therapies in clinical settings, due to their low immunogenic potential and their low oncogenic risk from host-genome integration and long-term gene expression in tissues and organs^46^. We selected muscle-tropic AAV serotype 2/9 vectors as a mouse delivery method for our Fn-REDLIP system. We deployed our approach in two separate AAV viral vectors: the constitutively expressed RL-responsive sensor Gal4-FnBphP (2571 bp) or Gal4-PnBphP (2289 bp) and the *trans-*activator LDB3-p65-HSF1 (1542 bp). The mice were intramuscularly injected in the left gastrocnemius muscle with a mixture of these two AAVs and the RL-responsive luciferase reporter AAV vector. Two weeks after AAV transduction **(Fig. 4d and Supplementary Fig. 7a)**, we illuminated the mice at an intensity of 20 mW/cm^2^ for 30 minutes once every three or four weeks. Bioluminescence imaging at different time points showed that the mice given this AAV-packaged Fn-REDLIP or Pn-REDLIP system with RL illumination can induce luciferase reporter expression for up to 3.5 months **(Fig. 4e, f, and Supplementary Fig. 8)**. The AAV2/9_Luc_ group (reporter only) had significantly decreased luciferase expression levels compared to the AAV2/9_Pn-Luc_-dark group; this apparent leakage was marginal when considered against the strong induction upon RL illumination **(Supplementary Fig. 7b, c)**. Collectively, these results indicate that the AAV-packaged Fn-REDLIP and Pn-REDLIP systems enable non-invasive and long-term regulation of gene expression *in vivo*.

### Optogenetic control of glucose homeostasis using an AAV-delivered Fn-REDLIP system in type 1 diabetic mice

We tested the Fn-REDLIP system for the on-demand production of therapeutic agents *in vivo* by deploying our approach in two separate AAV-2/9 vectors: the constitutively expressed RL-responsive sensor Gal4-FnBphP and a single vector which concatenated the constructs for the hybrid *trans-*activator module LDB3-p65-HSF1 and the P_RL_-driven insulin expression module. We then transduced type 1 diabetic (T1D) mice with a mixture of these two AAVs by intramuscular injection in the left gastrocnemius muscle (AAV2/9_INS_), and two weeks after AAV transduction, mice were illuminated with RL (660 nm, 20 mW/cm^2^) for 30 minutes twice a day or kept in the dark **(Fig. 5a, b)**. RL illumination of the Fn-REDLIP AAV-transduced mice (AAV2/9_INS_-660 nm) resulted in significantly elevated insulin levels compared to dark control AAV-transduced mice (AAV2/9_INS_-Dark) and to the STZ-induced T1D control mice (STZ) **(Fig. 5c)**. Moreover, the RL-induced insulin reduced blood glucose levels for up to 12 weeks, while the STZ group and AAV2/9_INS_-Dark group had continuously high blood glucose levels **(Fig. 5d)**. An intraperitoneal glucose tolerance test (IPGTT) showed a significant improvement in glucose tolerance in the AAV2/9_INS_-660 nm group **(Fig. 5e, f)**. These results demonstrate that the AAV-delivered Fn-REDLIP system regulates insulin expression to achieve long-term blood glucose control in T1D mice.

**Fig. 5.**
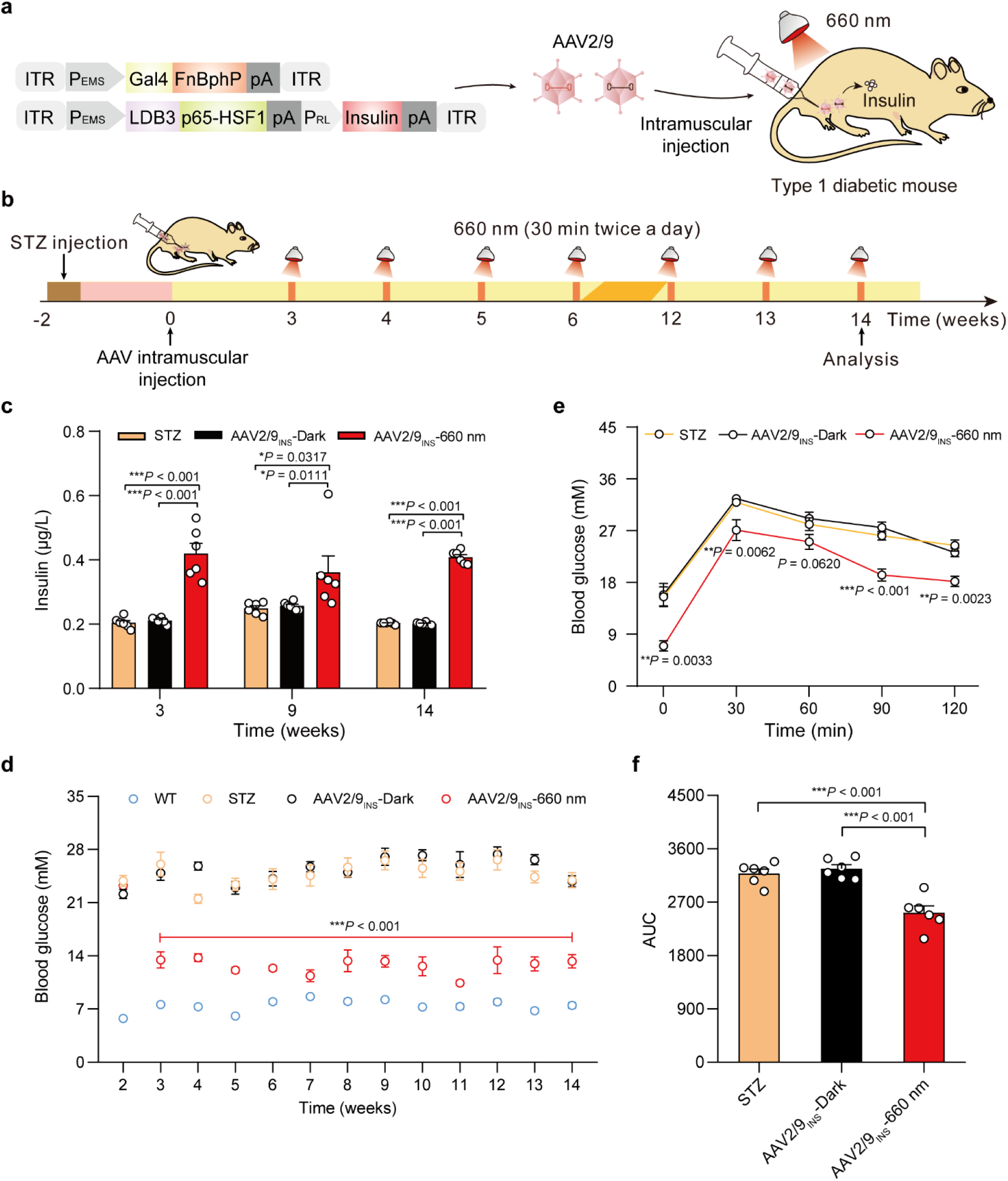
An AAV-delivered Fn-REDLIP system for long-term insulin expression to control blood glucose homeostasis in T1D mice. **a,** Schematic representation of the genetic configuration of the AAV2/9 vectors for the Fn-REDLIP system used in the T1D model mice experiments. **b,** Schematic representation of the experimental procedure and time schedule for AAV-delivered Fn-REDLIP-mediated insulin expression in T1D mice. T1D mice were intramuscularly injected in the left gastrocnemius muscle with a mixture of AAVs encoding the Fn-REDLIP system iteration containing pQL382 (ITR-P_EMS_-Gal4-FnBphP-pA-ITR) and the concatenated vector pQL388 (ITR-P_EMS_-LDB3-p65-HSF1-pA::P_RL_-EGFP-2A-insulin-pA-ITR) at an AAV titer of 2×10^11^ vg. After 2 weeks, the injected T1D mice were illuminated with or without RL at an intensity of 20 mW/cm^2^ for 30 minutes twice a day at the indicated time point (AAV2/9_INS_-660 nm or AAV2/9_INS_-Dark). The examined controls included non-model wild-type control mice (WT group) and the STZ-induced T1D control mice (STZ group). **c, d,** The AAV-delivered Fn-REDLIP system for long-term blood glucose control in T1D mice. T1D mice were intramuscularly injected as described in (B), followed by illumination with RL (660 nm, 20 mW/cm^2^, 30 minutes) twice daily. Blood insulin (c) was profiled weekly using a mouse insulin ELISA kit, and blood glucose (d) was profiled once weekly using a blood glucose meter. **e,** An intraperitoneal glucose tolerance test (IPGTT) was conducted at week 14 after AAV injection. **f,** Area under the curve (AUC) analysis of the IPGTT data in e. Data in c-f are presented as means ± SEM (*n* = 6 mice). *P* values were calculated by one-way ANOVA with multiple comparisons. *P* values in d, e were calculated by comparing the AAV2/9_INS_-Dark group with the AAV2/9_INS_-660 nm group.

We next assessed the capacity for turning insulin expression on and off *in vivo*. T1D mice were intravenously injected with a mixture of AAV-2/8 vectors (targeting the livers) encoding the Fn-REDLIP system (**Supplementary Fig. 9a**). After 2 weeks, the injected T1D mice were randomly divided into three groups: the dark control group (Dark), RL illumination group (660 nm), RL and FRL illumination group (660 nm+780 nm) (**Supplementary Fig. 9b, c)**. Compared to the dark control mice, the mice in RL illumination group received RL (660 nm, 20 mW/cm^2^) for 30 minutes twice a day (**Supplementary Fig. 9b**), while mice in RL and FRL group were first exposed to RL and transitioned to FRL (780 nm, 20 mW/cm^2^) at the indicated time (**Supplementary Fig. 9c**). After 24 hours of exposure, both RL and the combination of RL/FRL induced a significant increase in circulating insulin levels, resulting in a dramatic reduction in blood glucose levels. In contrast, the dark control mice exhibited no change in insulin and blood glucose levels (**Supplementary Fig. 9d, e**), indicating that the change in insulin and blood glucose depends entirely on RL and RL/FRL exposure. Moreover, after a subsequent 48-hour interval, the insulin and blood glucose returned to the same levels as in the dark control mice (**Supplementary Fig. 9d, e**), indicating the tunability of this system. Further, a second round of RL and RL/FRL exposure efficiently induced the increase of insulin and reduction of blood glucose levels in the RL and RL/FRL group (**Supplementary Fig. 9d, e**). These results indicate that this response could be turned on/off by adjusting the illumination exposure to achieve the desired insulin and blood glucose levels in T1D mice.

### Optogenetic control of TSLP expression using an AAV-delivered Fn-REDLIP system in high-fat diet (HFD)-induced obesity model mice

We next developed an Fn-REDLIP system for an *in vivo* gene therapy application to control the expression of thymic stromal lymphopoietin (TSLP), which has been recently reported to protect against obesity and obesity-related complications^43^. However, the overexpression of TSLP can cause airway inflammation and hyperreactivity^47, 48^. We constructed a relatively simple system comprising two AAV-2/9 vectors encoding the constitutively expressed RL-responsive sensor Gal4-FnBphP and the concatenated constructs for the hybrid *trans-*activator LDB3-p65-HSF1 and the P_RL_-driven TSLP expression module (**Fig. 6a**). We intramuscularly injected HFD-induced obesity model mice in the left gastrocnemius muscle with these two AAVs or two control AAVs containing RL-responsive sensor Gal4-FnBphP and the concatenated hybrid *trans-*activator vector LDB3-p65-HSF1 and the P_RL_-driven EGFP expression module (AAV2/9_TSLP_ or AAV2/9_EGFP_). After two weeks, the injected HFD mice were illuminated at an intensity of 20 mW/ cm^2^ for 30 minutes once every three days until week 6 (AAV2/9_TSLP_-660 nm) or were kept in the dark (AAV2/9_TSLP_-Dark) (**Fig. 6b**). AAV2/9_TSLP_-660 nm group had significantly increased TSLP levels compared to AAV2/9_TSLP_-Dark group, to AAV2/9_EGFP_-660 nm group, and untreated HFD control mice (**Fig. 6c**). Strikingly, after receiving the RL illumination for two weeks, the AAV2/9_TSLP_-660 nm group had significantly reduced body weight compared to the AAV2/9_TSLP_-Dark group, the AAV2/9_EGFP_-660 nm group, and the untreated HFD control mice; indeed, these mice had body weights that did not differ from wild type control mice at 8 weeks after the initial AAV transduction followed by illumination (**Fig. 6d, e**). Moreover, this treatment group displayed significantly decreased weights of adipose tissues (including epididymal white adipose tissue (eWAT), subcutaneous inguinal white adipose tissue (iWAT), and brown adipose tissue (BAT)) compared to the AAV2/9_TSLP_-Dark group, WT control mice, and untreated HFD control mice (**Fig. 6f**). A battery of metabolic tests, including a fasting blood glucose test (**Fig. 6g**), an intraperitoneal glucose tolerance test (IPGTT) (**Fig. 6h**), an insulin tolerance test (ITT) (**Fig. 6i**), a homeostatic model assessment of insulin resistance (HOMA-IR) test (**Fig. 6j**), as well as measuring serum and liver triglycerides (TGs) (**Fig. 6k, l**), consistently showed significant decreases in the AAV2/9_TSLP_-660 nm group compared to the relevant controls; these parameters did not differ from wild type control mice 8 weeks after the initial AAV transduction followed by illumination.

**Fig. 6.**
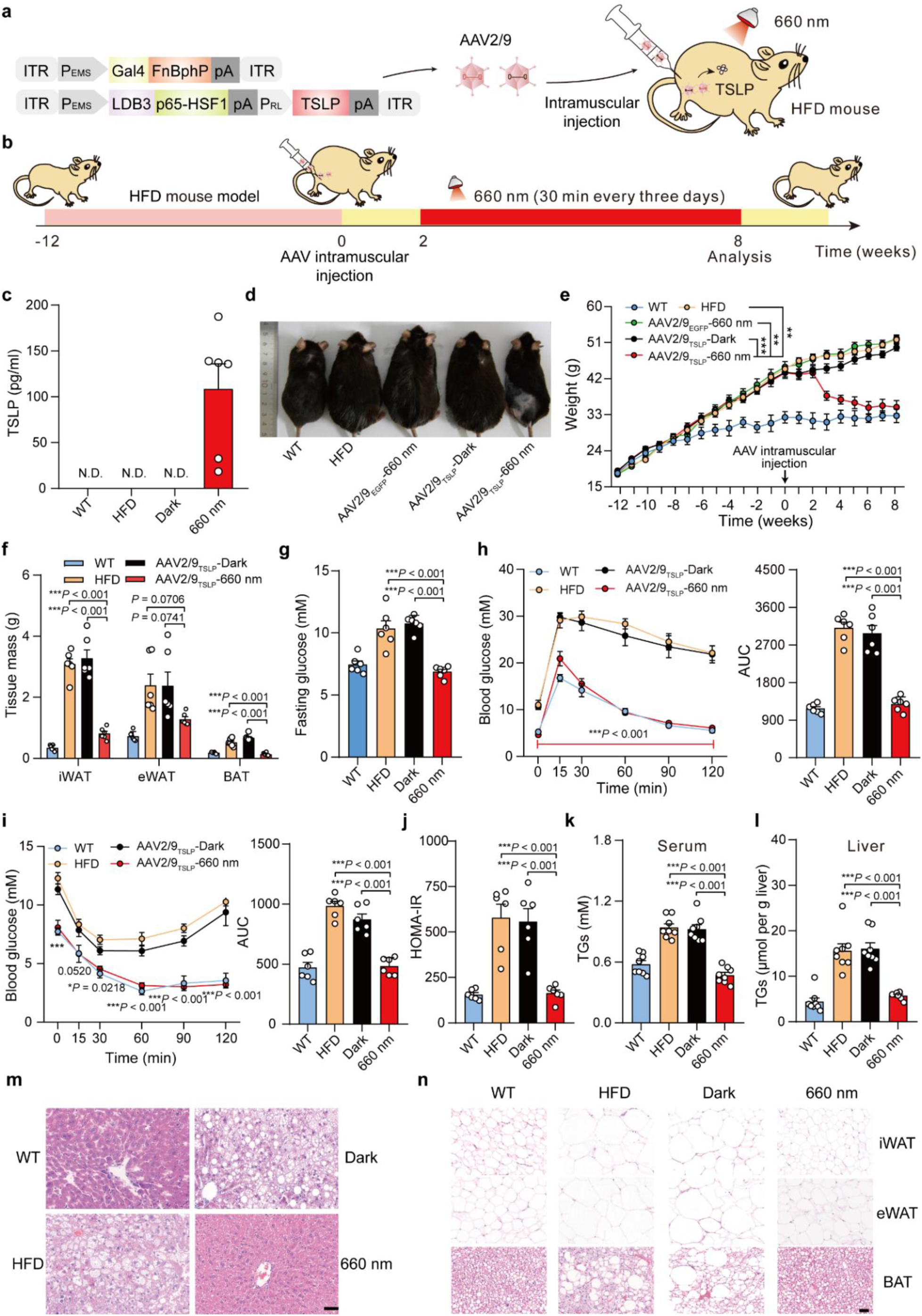
An AAV-delivered Fn-REDLIP system for long-term TSLP expression to control body weight in HFD mice. **a,** Schematic representation of the genetic configuration of the AAV2/9 vectors for the Fn-REDLIP system used for the HFD obesity model mice experiment. **b,** Schematic representation of the experimental procedure and time schedule for AAV-delivered Fn-REDLIP-mediated TSLP transgene expression in HFD mice. HFD mice were intramuscularly injected in the left gastrocnemius muscle with a mixture of AAVs encoding the Fn-REDLIP system iteration containing pQL382 (ITR-P_EMS_-Gal4-FnBphP-pA-ITR) and the concatenated vector pQL383 (ITR-P_EMS_-LDB3-p65-HSF1-pA::P_RL_-TSLP-pA-ITR) at an AAV titer of 2×10^11^ vg. After 2 weeks, the injected HFD mice were illuminated with or without RL illumination at an intensity of 20 mW/cm^2^ for 30 minutes every three days for 6 weeks (AAV2/9_TSLP_-660 nm or AAV2/9_TSLP_-Dark). The examined controls included non-model wild-type control mice (WT group), HFD control mice (HFD group), and HFD mice transduced with control EGFP AAVs (AAV-delivered Fn-REDLIP-mediated EGFP transgene expression) with RL illumination (AAV2/9_EGFP_-660 nm). **c,** The levels of TSLP expression in serum were quantified using a mouse TSLP ELISA kit. N.D., Not detected. **d,** Representative images of the AAV2/9_TSLP_-660 nm group at 8 weeks. The examined controls included the AAV2/9_TSLP_-Dark group, WT group, HFD group, and AAV2/9_EGFP_-660 nm group. **e-l,** Analysis of metabolic parameters of HFD mice treated with the AAV-delivered Fn-REDLIP system, including (e) Body weight, \*\*\**P* < 0.001, \*\**P* = 0.0011, \*\**P* = 0.0024; (f) adipose tissue (eWAT, iWAT, and BAT) weights; (g) Fasting blood glucose; (h) IPGTT (left) and AUC analysis of the IPGTT data (right); (i) ITT (left) and AUC analysis of the ITT data (right); (j) HOMA-IR; (k) Serum TG, and (l) Liver TG. **m, n**, Representative H&E staining of the liver (m) and adipose (n) tissues. Scale bars = 50 µm. Data in c and e-l are presented as means ±SEM (*n* = 6-8 mice). *P* values were calculated by one-way ANOVA with multiple comparisons. *P* values in h, i were calculated by comparing the AAV2/9_TSLP_-Dark group with the AAV2/9_TSLP_-660 nm group.

Hematoxylin and eosin staining revealed that the AAV2/9_TSLP_-660 nm group exhibited significantly smaller adipose droplets, decreased severity of hepatic steatosis in the liver (**Fig. 6m**), and smaller adipocyte sizes in adipose tissues (epididymal white adipose tissue (eWAT), inguinal white adipose tissue (iWAT), and brown adipose tissue (BAT)] (**Fig. 6n**). Notably, long-term observations of mice given this AAV-delivered Fn-REDLIP system showed no obvious adverse effects, including blood biochemistry [alanine aminotransferase (ALT), aspartate aminotransferase (AST), blood urea nitrogen (BUN), creatinine (CRE)], hematology [albumin/globulin ratio (A/G), counts for red blood cells (RBC), hemoglobin (HGB), platelet (PLT), total white blood cells (WBC), percentage of blood lymphocytes (LYMPH), blood monocytes (MONO), blood eosinophil (EO)], serum inflammatory cytokine (serum IL-6 and serum TNF-α), and immunoglobulin (serum IgG) (**Supplementary Figs. 10, 11**). These results demonstrate that the AAV-delivered Fn-REDLIP system enables precise optogenetic control of the therapeutic protein to implement effective gene therapy in animals.

We also achieved dosage control of TSLP with Fn-REDLIP in mouse livers. Briefly, intravenously injected HFD mice with a mixture of AAV2/8_TSLP_ vectors and exposed mice to different RL (20 mW/cm^2^) illumination time (0, 5, 30 minutes) every three days for 4 weeks (**Supplementary Fig. 12a, b**). The mice of the 30 minutes exposure group had a significantly higher TSLP levels. They showed the expected attendant reduction in body weight (**Supplementary Fig. 12c**-e). These results demonstrate that the AAV-delivered Fn-REDLIP system enables dose-dependent regulation of transgene expression through varying durations of light exposure in deep tissues of mice.

## DISCUSSION

Optogenetics allows for the traceless and remote control of living cells, tissues, and organisms with tailored functions. Moreover, optogenetics is moving from basic research toward therapeutic applications ^14, 49–51^. In this study, we have successfully developed the REDLIP system (Fn-REDLIP and Pn-REDLIP) based on the chimeric photosensitive protein BphPs and their binding partner LDB3, which enables control of gene expression in a light-intensity and light-duration dependent manner, as well as provides reversible features in response to light. The Fn-REDLIP system components can be delivered by AAV to control *in vivo* transgene expression.

Compared with existing red/far-red light-responsive optogenetic systems, the REDLIP activation requires a shorter illumination time (10 seconds of exposure to RL), fast ON/OFF kinetics (10 seconds ON and 1 minute OFF), and strong activation of gene expression (> 100-fold). Moreover, our system does not require users to introduce exogenous chromophore, whereas other red/far-red optogenetic systems based on PhyB /PIF6, PhyA/FHY1, or PhyA/FHL each require users to supply an exogenous pigment cofactor^24, 30^ or produce a cofactor through co-expression of biosynthesis genes^32, 52^. In addition, the Fn-REDLIP_cas_ and Pn-REDLIP_cas_ systems activation also requires a shorter illumination time (10 seconds) and minutes (30 minutes) in mammalian cells (> 326-fold) and mice (> 56-fold), respectively, while the recently reported Red-CPTS^38^ requires repetitive pulses of illumination over 24 hours. We also anticipate that the REDLIP system can be adaptable to the Cre-*loxP* system to achieve red light-activatable Cre-mediated DNA recombination, which can be used for gene insertion, deletion, and inversion in various animal models and the creation of transgenic mouse models.

Translational optogenetics would benefit from a safe and effective gene delivery method. AAVs have recently gained popularity as gene delivery vehicles with acceptable safety, low immunogenicity, and long-term gene expression potential^53^. The small size of the Fn-REDLIP system enables it to be packaged into AAVs to achieve RL-inducible protein expression. We have demonstrated that the Fn-REDLIP system can be delivered to the skeletal muscles/livers of mice by AAV and enables the light-regulated expression of insulin in diabetes model mice to reduce blood glucose or expression of TSLP in HFD mice to control body weight. The REDLIP system is well-suited to treat diseases that require long-term yet controllable expression of therapeutic proteins.

There is still room for improvement with regard to the delivery method of our system. Developments in capsid design for AAVs variants^54–56^ and non-viral vectors^22, 57, 58^ can advance gene delivery strategies, which could help facilitate the delivery of our system into targeted tissues. Moreover, more compact systems and existing or new miniature engineered photoreceptors will be developed to simplify their applications *in vivo*. To avoid potential gene expression leakiness induced by ambient light, REDLIP-mediated gene therapy could be administered via subcutaneous muscle delivery using an LED patch attached to the skin, effectively shielding it from ambient light exposure^59^. The versatility of our system should enable it to control various protein-based drugs, including enzymes, peptide hormones, and antibodies. For instance, the REDLIP system could be used to induce the expression of urate oxidase to treat gout^5^ or the expression of parathyroid hormone (PTH) to treat hyperparathyroidism^60^.

Collectively, the REDLIP system represents a new optogenetic tool allowing for efficient gene expression without exogenous administration of a chromophore in mammals in a timescale of seconds with the capacity for deep penetration and invasiveness. We expect that our AAV-delivered REDLIP system will accelerate the progression of optogenetic therapies toward clinical applications.

## MATERIALS AND METHODS

### Ethical statement

The animal experiments were performed according to the protocol approved by the East China Normal University (ECNU) Animal Care and Use Committee. They were in accordance with the Ministry of Science and Technology of the People’s Republic of China on Animal Care Guidelines. The protocol was approved by the ECNU Animal Care and Use Committee (protocol ID: m20220505, R+RB20210101). All mice were euthanized after the termination of the experiments.

### Plasmid construction

Construction details of the plasmids are provided in **Supplementary Table 1**. The DNA sequences for REDLIP components in this study are listed in **Supplementary Table 6**. The plasmids were constructed by Gibson assembly according to the manufacturer’s instruction (MultiS One Step Cloning Kit, Catalog no. C113-01, Vazyme). DNA sequencing (Shanghai Saiheng Biotechnology) confirmed the sequences.

### Cell culture and transfection

Human embryonic kidney cells (HEK-293T, CRL-11268, ATCC), telomerase-immortalized human mesenchymal stem cells (hMSC-TERT), human cervical adenocarcinoma cells (HeLa, CCL-2, ATCC), HEK-293-derived Hana3A cells engineered for constitutive expression of RTP1, RTP2, REEP1, and G_αoλϕ_^61^; mouse chondroblast cells (ATDC5, CRL-3419, ATCC) were cultured in Dulbecco’s modified Eagle’s medium (DMEM, Catalog no. 12100061, Gibco) supplemented with 10% (v/v) fetal bovine serum (Catalog no. FBSSA500-S, AusGeneX), and 1% (v/v) penicillin/streptomycin solution (Catalog. no. ST488-1/ST488-2, Beyotime). All the cell lines were cultured at 37 ℃ in a humidified atmosphere containing 5% CO_2_, and were regularly tested for the absence of *Mycoplasma* and bacterial contamination. The concentration and viability of the cell lines were evaluated using a Countess II Automated cell counter (AMEP4746, Life Technologies).

All cells except for ATDC5 were transfected using an optimized polyethyleneimine (PEI)-based protocol. Briefly, 6×10^4^ cells per well were plated into a 24-well plate and cultured for 18 hours. The cells were subsequently co-transfected with corresponding plasmid mixtures for 6 hours with 50 μL PEI (Catalog no. 24765, Polysciences; molecular weight 40,000, stock solution 1 mg/mL in ddH_2_O; PEI and DNA at a mass ratio of 3:1 for HEK-293T, hMSC-TERT, and Hana3A and at a mass ratio of 5:1 for HeLa.). For the ATDC5 cells, 6×10^4^ cells per well were plated into a 24-well plate, cultured for 18 hours, and co-transfected with corresponding plasmid mixtures using Lipofectamine™ 3000 (Catalog. no. L3000015, Thermo Fisher Scientific) according to the manufacturer’s instructions.

### The ON/OFF performance of the Pn-REDLIP and Fn-REDLIP systems in HEK-293T cells

HEK-293T cells were plated into a 24-well plate (6×10^4^ per well), cultivated to 70-90% confluence, and co-transfected with a total of 375 ng of the plasmids encoding the Pn-REDLIP system [pNX12, pQL325, and pYZ430 at a 2:2:1 (w/w/w) ratio] or a total of 375 ng of plasmids encoding the Fn-REDLIP system [pNX12, pQL326, and pYZ430 at a 2:2:1 (w/w/w) ratio]. Subsequently, the cells were illuminated with RL (660 nm, 1.0 mW/cm^2^) for 3 seconds (ON) or kept in the dark (OFF). The culture medium was refreshed every 24 hours under green light (530 nm) with concomitant reversal of illumination conditions, and SEAP production was quantified every 12 hours for 72 hours.

### Animals

All animals were approved by the Institutional Animal Care and Use Committee of Shanghai and conducted in accordance with the National Research Council Guide for Care and Use of Laboratory Animals. The experimental animals included 4- or 6-week-old or body weight ∼25 g C57BL/6 male mice. Mice were reared in East China Normal University Laboratory Animal Center and kept with a standard alternating 12-hour light/12-hour dark cycle during the construction of T1D and HFD mouse models and before AAV transduction. After AAV transduction, the mice in the dark group were consistently maintained in darkness except for specific procedures such as blood sampling, changing water bottles, and cleaning cages, which were conducted under green light (530 nm).

### The REDLIP-mediated transgene expression in mammalian cells and mice

For REDLIP-mediated transgene expression in mammalian cells, 6×10^4^ cells were plated in a 24-well plate and cultivated to 70-90% confluency during transfection. For the Pn-REDLIP system, the cells were co-transfected with a total of 375 ng of the Pn-REDLIP system-encoding plasmids [pNX12 (P_hCMV_-2NLS-LDB3-p65-HSF1-pA), pYZ430 (P_RL_-SEAP-pA) and pQL325 (P_hCMV_-Gal4-PnBphP-pA) at a 2:1:2 (w/w/w) ratio]. For the Fn-REDLIP system, the cells were co-transfected with a total of 375 ng of the Fn-REDLIP system-encoding plasmids [pNX12, pYZ430, and pQL326 (P_hCMV_-Gal4-FnBphP-pA) at a 2:1:2 (w/w/w) ratio]. Six hours after transfection, the cell culture medium was replaced with a fresh medium. Twenty-four hours after transfection, cell culture plates were placed below a custom-designed 4 ×6 red light LED array (660 nm; Shenzhen Bested Opto-electronic; each LED centered above a single well), and the cells were illuminated with RL (660 nm, 2.0 mW/cm^2^, 10 seconds, unless explicitly indicated). The light intensity was determined by an 8230E optical power meter (ADC Corporation). SEAP production was quantified 24 hours after illumination. The ambient light intensity (∼750 Lux) was determined by a digital illuminometer (DL333204, Ningbo Deli Tools Co., Ltd.).

For REDLIP-mediated transgene expression in mice, the mice (C57BL/6, male, 6-week-old) were hydrodynamically injected with 2 mL (10% of the body weight in grams) Ringer’s solution (0.147 M NaCl, 1.13 M CaCl_2_, 4×10^−3^ M KCl) containing plasmids encoding the Dr-REDLIP, Pn-REDLIP, or Fn-REDLIP. Each mouse was hydrodynamically injected with 375 μg of the plasmid DNA encoding the Dr-REDLIP [pQL236 (P_hCMV_-3NLS-LDB3-p65-HSF1-pA), pQL217(P_hCMV_-Gal4-DrBphP-pA), and pYZ450 (P_RL_-Luciferase-pA) at a 2:2:1 (w/w/w) ratio] or a total of 375 μg of the Pn-REDLIP [pQL236, pQL325, and pYZ450 at a 2:2:1 (w/w/w) ratio] or a total of 375 μg of the Fn-REDLIP [pQL236, pQL326, and pYZ450 at a 2:2:1 (w/w/w) ratio]. Sixteen hours after plasmid injection, mice were illuminated with RL (660 nm LED, 20 mW/cm^2^) for different time periods (0 to 2 hours). Eight hours after initial illumination, bioluminescence images of the mice were obtained using an *IVIS* Lumina II *in vivo* imaging system (Perkin Elmer), and analyzed with the Living Image software (version 4.3.1).

### *In vivo* bioluminescence and imaging

Each mouse was intraperitoneally injected with 15 mg/mL D-Luciferin solution (150 mg/kg; Catalog no. luc001, Shanghai Sciencelight Biology Science & Technology) and anesthetized with 2% isoflurane (Catalog no. R510-22-10, RWD Life Science) dissolved in oxygen using an economical animal anesthesia machine (HSIV-S, Raymain). Ten minutes after Luciferin injection, bioluminescence images of the mice were taken using the IVIS Lumina II *in vivo* imaging system (Perkin Elmer). Regions of interest were quantified using the Living Image software (version 4.3.1).

### The Pn-REDLIP_cas_ and Fn-REDLIP_cas_ systems for activation of endogenous gene transcription in mammalian cells and mice

For REDLIP_cas_-mediated endogenous gene transcription in mammalian cells, the different mammalian cells (HEK-293T, hMSC-TERT, HeLa, and Hana3A) were co-transfected with a total of 510 ng of the plasmids encoding the Pn-REDLIP_cas_ [pNX12, pQL325, pDQ100 (P_RL_-MS2-p65-HSF1-pA), pSZ69 (P_hCMV_-dCas9-pA), and the corresponding two sgRNAs (**Supplementary Table 2**) at a 15:15:1:10:5:5 (w/w/w/w/w/w) ratio] or a total of 510 ng of the plasmids encoding the Fn-REDLIP_cas_ [pNX12, pQL326, pDQ100, pSZ69, and the corresponding two sgRNAs (**Supplementary Table 2**) at a 15:15:1:10:5:5 (w/w/w/w/w/w) ratio]. The negative control cells were transfected with the empty pcDNA3.1(+) vector. Twenty-four hours after transfection, the cells were illuminated for different time periods (660 nm, 0-10 seconds) and light intensities (660 nm, 0 to 2.0 mW/cm^2^). Cells were then collected, and total RNA was extracted for qPCR analysis 24 hours after illumination.

For REDLIP_cas_-mediated endogenous gene transcription in mice, the mice (C57BL/6, male, 6-week-old) were hydrodynamically injected with 2 mL (10% of the body weight in grams) Ringer’s solution containing a total of 410 μg of the plasmid DNA encoding the Fn-REDLIP_cas_ [pQL236, pQL326, pYZ561 (P_U6_-sgRNA1*_Ascl1_*-pA::P_U6_-sgRNA2*_Ascl1_*-pA::P_hCMV_-dCas9-pA), and pDQ100 at a 15:15:10:1 (w/w/w/w) ratio] via tail vein. Sixteen hours after plasmid injection, mice were illuminated with RL (660 nm LED, 20 mW/cm^2^, 30 minutes). Negative control mice were hydrodynamically injected with Ringer’s solution. At 16 hours following illumination, the mice were sacrificed, and liver tissues were collected for qPCR analysis of *Ascl1* expression. qPCR primers used in this study are listed in **Supplementary Table 3**. All samples were normalized to housekeeping gene *glyceraldehyde 3-phosphate dehydrogenase* (*Gapdh*) values, and the results are expressed as the relative mRNA level normalized to that in the negative control using the standard 2^−ΔΔCT^ method.

### Construction of mouse model of type 1 diabetes (T1D)

The type 1 diabetes mouse model (T1D) was induced by streptozotocin (STZ) injection. Briefly, adult male mice (C57BL/6, with body weight ∼25 g) were fasted for 16 hours and intraperitoneally injected with STZ (Catalog no. S0130, Merck; 50 mg/kg in 0.1 M citrate buffer, pH 4) every day for five days. Two weeks after the initial injection, fasted mice with hyperglycemia over 16.7 mmol/L glucose were considered diabetic and used for further animal experiments.

### AAV production

For the AAV-delivered Fn-REDLIP/Pn-REDLIP system by intramuscular injection, the adeno-associated viral vector (serotype 2/9) was selected to package the following: RL-responsive sensor Gal4-FnBphP or Gal4-PnBphP expression vector (pQL382, ITR-P_EMS_-Gal4-FnBphP-pA-ITR; pNX257, ITR-P_EMS_-Gal4-PnBphP-pA-ITR), the hybrid *trans-*activator LDB3-p65-HSF1 expression vector (pNX137, ITR-P_EMS_-LDB3-p65-HSF1-pA-ITR), the light-inducible luciferase reporter pQL271 (ITR-P_RL_-Luciferase-pA-ITR), the single vector concatenated the constructs for the expression of the *trans-*activator LDB3-p65-HSF1 module, and the P_RL_-driven insulin expression module (pQL388, ITR-P_EMS_-LDB3-p65-HSF1-pA::P_RL_-EGFP-2A-insulin-pA-ITR), the P_RL_-driven TSLP expression module (pQL383, ITR-P_EMS_-LDB3-p65-HSF1-pA::P_RL_-TSLP-pA-ITR), or the P_RL_-driven EGFP expression module (pNX221, ITR-P_EMS_-LDB3-p65-HSF1-pA::P_RL_-EGFP-pA-ITR). These AAVs were produced by Shanghai Taitool Bioscience, purified AAVs were titrated using quantitative PCR and concentrated in PBS to ∼2×10^13^ vector genomes (vg) per mL.

For AAV-delivered Fn-REDLIP system by intravenous injection, the adeno-associated viral vector (serotype 2/8, targeting the livers) was selected to package the following: RL-responsive sensor Gal4-FnBphP expression vector (pNX177, ITR-P_hCMV_-Gal4-FnBphP-pA-ITR), the single vector concatenated the constructs for the expression of the *trans-*activator LDB3-p65-HSF1 module, the P_RL_-driven insulin expression module (pQL318, ITR-P_hCMV_-LDB3-p65-HSF1-pA::P_RL_-EGFP-2A-insulin-pA-ITR), and the P_RL_-driven TSLP expression module (pNX166, ITR-P_hCMV_-LDB3-p65-HSF1-pA::P_RL_-TSLP-pA-ITR). These AAVs were produced by Shanghai Taitool Bioscience, purified AAVs were titrated using quantitative PCR and concentrated in PBS to ∼2×10^13^ vector genomes (vg) per mL.

### Optogenetic control of luciferase expression using AAV-delivered Fn-REDLIP/Pn-REDLIP system in mice

Wild-type C57BL/6 mice (6-week-old) were randomly divided into two groups and were intramuscularly injected into the left gastrocnemius muscle using an Omnican® 40 (0.3 mm × 8 mm) syringe with a mixture of 50 μL AAVs (serotype 2/9) containing three AAV vectors: the Gal4-FnBphP (pQL382, ITR-P_EMS_-Gal4-FnBphP-pA-ITR, 2×10^11^ vg) or Gal4-PnBphP (pNX257, ITR-P_EMS_-Gal4-PnBphP-pA-ITR, 2×10^11^ vg) expression vector, the hybrid *trans-*activator LDB3-p65-HSF1 expression vector (pNX137, ITR-P_EMS_-LDB3-p65-HSF1-pA-ITR, 2×10^11^ vg) and the luciferase reporter pQL271 (ITR-P_RL_-Luciferase-pA-ITR, 2×10^11^ vg). The control group received an intramuscular injection (left gastrocnemius muscle) of the RL-responsive luciferase reporter AAV vector (AAV2/9_luc_, 2×10^11^ vg). Two weeks after injection, the mice were illuminated with red light at an intensity of 20 mW/cm^2^ for 30 minutes once every three or four weeks. Eight hours after illumination, bioluminescence images of the mice were obtained using an *IVIS* Lumina II *in vivo* imaging system (Perkin Elmer), and analyzed with the Living Image software (version 4.3.1).

### Optogenetic control of blood glucose homeostasis using an AAV-delivered Fn-REDLIP system in T1D mice

For AAV-delivered Fn-REDLIP system controlling intramuscular insulin expression experiment, T1D mice were intramuscularly injected in the left gastrocnemius muscle with a mixture of 50 μL AAVs (serotype 2/9) packaging two AAV vectors: the Gal4-FnBphP expression vector (pQL382, ITR-P_EMS_-Gal4-FnBphP-pA-ITR, 2×10^11^ vg) and a single vector which concatenated the constructs for the hybrid *trans-*activator LDB3-p65-HSF1 module and the P_RL_-driven insulin (pQL388, ITR-P_EMS_-LDB3-p65-HSF1-pA::P_RL_-EGFP-2A-insulin-pA-ITR, 2×10^11^vg) expression module. Two weeks after transduction, the injected T1D mice were illuminated with or without red light at an intensity of 20 mW/cm^2^ for 30 minutes twice a day for 12 weeks. Twenty-four hours after first illumination, blood insulin was profiled in the 3^rd^, 9^th^, and 14^th^ week using a mouse insulin ELISA kit, and blood glucose was profiled once a week using a blood glucose meter. The intraperitoneal glucose tolerance test (IGTT) was performed in the 14^th^ week after the AAV injection.

For the AAV-delivered Fn-REDLIP system controlling hepatic insulin expression experiment, T1D mice were intravenously injected with a mixture of 50 μL AAVs (serotype 2/8) packaging two AAV vectors: the Gal4-FnBphP expression vector (pNX177, ITR-P_hCMV_-Gal4-FnBphP-pA-ITR, 2×10^11^ vg) and the single vector concatenated the constructs for the expression of the *trans-* activator LDB3-p65-HSF1 module and the P_RL_-driven insulin expression module (pQL318, ITR-P_hCMV_-LDB3-p65-HSF1-pA::P_RL_-EGFP-2A-insulin-pA-ITR, 2×10^11^ vg). After 2 weeks, the injected T1D mice were randomly divided into three groups: i) Dark group: T1D mice kept in the dark; ii) 660 nm group: T1D mice exposed to RL (660 nm, 20 mW/cm^2^) for 30 minutes twice a day; iii) 660 nm+780 nm group: T1D mice exposed to RL (660 nm, 20 mW/cm^2^) for 30 minutes twice a day (ON), followed by exposure to FRL (780 nm, 20 mW/cm^2^) for 30 minutes twice a day (OFF) after a 3-hour interval. Each group of AAV-injected T1D mice underwent the procedure every 3 days for a total of 6 days. Starting 24 hours after the first RL illumination, blood insulin was profiled on days 1, 2, 4, 5, and 7 using a mouse insulin ELISA kit; blood glucose was profiled at the same time points using a blood glucose meter.

### Optogenetic control of TSLP expression using an AAV-delivered Fn-REDLIP system in obesity model mice

For the AAV-delivered Fn-REDLIP system controlling intramuscular TSLP expression experiment, the HFD mouse model was created by feeding a high-fat diet. The male mice (C57BL/6, 4-week-old) were fed high-fat diets consisting of 60 kcal% fat (Catalog no. D12492, Research Diets) for 13 weeks until the body weight exceeded 40 g. Subsequently, the HFD mice were intramuscularly injected in the left gastrocnemius muscle with a mixture of 50 μL AAVs (serotype 2/9) packaging two AAV vectors: the Gal4-FnBphP expression vector (pQL382, ITR-P_EMS_-Gal4-FnBphP-pA-ITR, 2×10^11^ vg) and the single vector concatenated the constructs for the expression of the *trans-*activator LDB3-p65-HSF1 module and the P_RL_-driven TSLP module (pQL383, ITR-P_EMS_-LDB3-p65-HSF1-pA::P_RL_-TSLP-pA-ITR, 2×10^11^ vg) or the P_RL_-driven EGFP module (pNX221, ITR-P_EMS_-LDB3-p65-HSF1::pA-P_RL_-EGFP-pA-ITR, 2×10^11^ vg). Two weeks after the AAV transduction, the injected HFD mice were illuminated with or without RL illumination at an intensity of 20 mW/cm^2^ for 30 minutes every three days for 6 weeks. The examined controls included non-model wild-type control mice, HFD control mice, and HFD mice transduced with control EGFP AAVs with RL illumination. Twenty-four hours after illumination, blood samples were collected from mouse retro-orbital sinus, transferred to ethylenediaminetetraacetic acid (EDTA) coated mini vacutainer tubes (Catalog no. BD-68784, BD Biosciences), and allowed to clot at 37°C for 30 minutes and then 4°C for 2 hours. The clotted blood was centrifuged at 1000 ×*g* for 10 minutes to get the serum, and serum TSLP levels were measured using a mouse TSLP ELISA Kit. Mice were weighed weekly, and the metabolic parameters were assessed, including fasting blood glucose, IPGTT, ITT, HOMA-IR, serum TG, and liver TG. The H&E staining of liver and adipose tissues was analyzed on the 8^th^ week after the AAV transduction.

For the AAV-delivered Fn-REDLIP system controlling hepatic TSLP expression experiment, HFD mice were intravenously injected with a mixture of 50 μL AAVs (serotype 2/8) packaging two AAV vectors: the Gal4-FnBphP expression vector (pNX177, ITR-P_hCMV_-Gal4-FnBphP-pA-ITR, 2×10^11^ vg) and the single vector concatenated the constructs for the expression of the *trans-* activator LDB3-p65-HSF1 module and the P_RL_-driven TSLP expression module (pNX166, ITR-P_hCMV_-LDB3-p65-HSF1-pA::P_RL_-TSLP-pA-ITR, 2×10^11^ vg). Two weeks after the AAV transduction, the injected HFD mice were illuminated with RL at an intensity of 20 mW/cm^2^ for 5 or 30 minutes every three days for 4 weeks. The examined controls include non-model wild-type control mice (WT group), untreated HFD control mice (HFD group), and the AAV-injected HFD mice without illumination (AAV2/8_TSLP_-Dark). Twenty-four hours after illumination, blood samples were collected, and serum TSLP levels were measured using a mouse TSLP ELISA Kit. All the mice were weighed weekly.

### Statistical analysis

Unless otherwise mentioned, all *in vitro* data represent means ±SD of three independent biological replicates. For the animal experiments, each treatment group consisted of randomly selected mice (*n* = 4-8). Comparisons between groups were performed using a two-tailed unpaired *t*-test or one-way ANOVA with multiple comparisons, and the results are expressed as means ± SEM. Differences were considered statistically significant at *\*P* < 0.05, very significant at \*\**P* < 0.01, and extremely significant at \*\*\**P* < 0.001. Neither animals nor samples were excluded from the study. GraphPad Prism 8 software was used for statistical analysis. *n* and *P* values are described in the figure legends.

## Acknowledgments

This work was financially supported by grants from the National Key R&D Program of China, Synthetic Biology Research (no. 2019YFA0904500), the National Natural Science Foundation of China (no. 31971346 and 31861143016), the Science and Technology Commission of Shanghai Municipality (no. 22N31900300 and 18JC1411000), and the Fundamental Research Funds for the Central Universities to H.Y. This work was also partially supported by National Key R&D Program of China (no. 2019YFA0110802), the National Natural Science Foundation of China (no. 32171414), the Natural Science Foundation of Shanghai (no. 23ZR1419500), and the Nature Science Foundation of Chongqing, China (no. CSTB2022NSCQ-MSX0461) to M.W. We also thank the support from the CAS Youth Interdisciplinary Team and the ECNU Multifunctional Platform for Innovation (011) for supporting the mice experiments and the Instruments Sharing Platform of the School of Life Sciences, ECNU.

## Author contributions

H.Y. conceived the project. H.Y., L.Q., M.W., and L.N. designed the experiments, analyzed the results, and wrote the manuscript. L.Q., L.N., Z.W., D.K., and G.Y. performed the experimental work. L.Q., M.W., L.N., and Z.W. designed, analyzed, and interpreted the experiments. All authors edited and approved the manuscript.

## Data and materials availability

All data associated with this study are presented in the paper or the Supplementary Information. All genetic components related to this paper are available with a material transfer agreement and can be requested from H.Y.

