## Supplementary Information for "A sensitive red/far-red photoswitch for controllable gene therapy in mouse models of metabolic diseases"

**Affiliations:**

### Materials and Methods

#### SEAP assay

A p-nitrophenylphosphate-based light absorbance time course assay was used to quantify the production of human placental SEAP. Briefly, 100 µL cell culture supernatants were heat-inactivated at 65°C for 30 minutes. 120 µL of substrate solution [100 µL of 2 × SEAP buffer (pH 9.8) containing 20 mM L-homoarginine hydrochloride (Catalog no. A602842, Sangon Biotech), 1 mM $MgCl_2$ (Catalog no. A610328, Sangon Biotech), 21% (w/w) diethanolamine (Catalog no. A600162, Sangon Biotech), and 20 µL of p-NPP substrate solution (Catalog no. 333338-18-4, Sangon Biotech) containing 120 mM p-nitrophenylphosphate] was added to 80 µL heat-inactivated supernatants. The time course of the absorbance at 405 nm was measured by the Synergy H1 hybrid multi-mode microplate reader (BioTek Instruments) with Gen5 software (version 2.04).

#### Spatial regulation of transgene expression and fluorescence imaging

HEK-293T cells were plated into a 10-cm dish ($3\times10^6$) and co-transfected a total of 12.5 µg of the plasmids encoding the Pn-REDLIP system [pNX12 ($P_{hCMV}$-2NLS-LDB3-p65-HSF1-pA), pQL325 ($P_{hCMV}$-Gal4-PnBphP-pA), and pDQ63($P_{RL}$-EGFP-pA) at a 2:2:1 (w/w/w) ratio] or a total of 12.5 µg of the plasmids encoding the Fn-REDLIP system [pNX12, pQL326 ($P_{hCMV}$-Gal4-FnBphP-pA), and pDQ63 at a 2:2:1 (w/w/w) ratio]. Twenty-four hours after transfection, the cells were placed on the display of a Vivo X9 smartphone with the pattern "PNTE" or "FNTE" and were illuminated with red light for 15 minutes (display brightness set to 100%, 40 µW/cm$^2$). Fluorescence images of EGFP expression were acquired 24 hours after illumination using ChemiScope 4300 Pro imaging equipment (Clinx).

#### qPCR analysis

According to the manufacturer's instructions, cells and tissue samples were harvested for total RNA isolation using an RNAiso Plus Kit (Catalog no. 9108, Takara). A total of 1 µg RNA was reverse transcribed into cDNA using a HiScript® II 1st Strand cDNA Synthesis Kit (+gDNA wiper) (Catalog no. R212-01, Vazyme) according to the manufacturer's instructions. qPCR was performed on a Real-Time PCR Instrument (LightCycler®96, Roche) using ChamQ Universal SYBR qPCR Master Mix (Catalog no. Q711-02, Vazyme) to detect the target genes. The amplification conditions of qPCR were as follows: 95°C for 10 minutes followed by 40 cycles at 95°C for 30 s, 55°C for 30 s, and 72°C for 30 s, and a final extension at 72°C for 10 min. The qPCR primers used in this study are listed in **table S3**. All samples were normalized to the human/mouse housekeeping gene *glyceraldehyde 3-phosphate* *dehydrogenase* (*Gapdh*), and the results were expressed as a relative mRNA expression level using the standard $2^{-\Delta\Delta CT}$ method.

**The enzyme-linked immunosorbent assay (ELISA)**

Insulin and TSLP in mouse serum were quantified using the mouse insulin ELISA Kit (Catalog no. 10-1247-01, Mercodia) and the mouse TSLP ELISA Kit (Catalog no. MTLP00, R&D), respectively. The levels of interleukin 6 (IL-6), and tumor necrosis factor α (TNF-α) were quantified using the mouse IL-6 ELISA Kit (Catalog no. EK206, Multi Sciences), and the mouse TNF-α ELISA Kit (Catalog no. EK282, Multi Sciences). IgG in mouse serum was quantified using the mouse IgG ELISA Kit (Catalog no. EK271, Multi Sciences). All samples for the ELISA assay were tested according to the manufacturer's instructions.

**Triglycerides measurement**

To measure triglycerides (TGs) in liver and serum, mouse liver and serum samples were collected 8 weeks after AAV injection. Liver tissues (20-100 mg) were homogenized with 9 μL absolute ethanol (Catalog no. A500737, Sangon Biotech) per mg liver tissue, smashed at 4°C using a low-temperature freezing grinding instrument (JXFSTPRP-CL, Thundersci), and centrifugated at 1000 × *g* for 10 minutes at 4°C. Blood was collected from the mouse eye socket and centrifugated at 1000 × *g* for 15 minutes to obtain serum. Then, the supernatants of liver tissues and serum TG levels were measured by a triglyceride assay kit (Catalog no. A110-2-1, Nanjing Jian Cheng Bioengineering Institute) according to the manufacturer's instructions.

**Intraperitoneal glucose tolerance test (IPGTT) in mice**

Mice were fasted overnight for 16 hours, and intraperitoneally injected with 10 μL 20% w/v D-glucose dissolved in 0.85% NaCl solution per g of body weight. Blood glucose levels were measured from tail vein blood samples at 0, 15, 30, 60, 90, and 120 minutes after D-glucose injection using a handheld glucometer (Exactive Easy III, MicroTech Medical). The 0-minute sample was used to determine the fasting plasma glucose level. The trapezoidal rule determined the area under the curve (AUC) for IPGTT.

**Insulin tolerance test (ITT) in mice**

Mice were fasted for 4 hours and intraperitoneally injected with 0.75 U/kg insulin solution (Catalog no.11061-68-0, Sigma). Blood samples were obtained via tail vein at 0, 15, 30, 60, 90, and 120 minutes after insulin injection, and the plasma insulin levels were determined by ELISA (Catalog no.10-1247-01, Mercodia) assay. The HOMA-IR (Homeostatic Model Assessment for Insulin Resistance) index was calculated using the standard formula: HOMA-IR index = fasting concentration of insulin (μIU/mL) × fasting concentration of glucose (mM)/22.5.

**Complete blood count and kidney function analysis**

Mice were euthanized, and their whole blood was collected and immediately analyzed for complete blood count using the Sysmex XT-2000iV hematology analyzer (Sysmex). The parameters of hepatic function include alanine aminotransferase (ALT), aspartate aminotransferase (AST), and albumin/globulin ratio (A/G). The parameters of kidney function, including creatinine (CRE) and blood urea nitrogen (BUN), were measured using an automatic biochemical analyzer BX-3010 (Sysmex). Plasma and serum samples were simultaneously analyzed for standard biochemical analytes.

**Hematoxylin and eosin (H&E) staining of liver and adipose tissues**

The liver and adipose tissue samples were collected and fixed in 4% paraformaldehyde (Catalog no. G1101, Servicebio) overnight at room temperature. The fixed samples were gently dehydrated by immersing them in a graded series of alcohol solutions, cleaned in xylene, embedded in paraffin, and sliced into 4 μm-thick sections with a rotary microtome (Leica RM2235, Manual Rotary Microtome). These sections were stained with an H&E Staining Kit (Catalog no. G1005, Servicebio) according to the manufacturer's instructions and were observed using an upright microscope (BX53, Olympus).

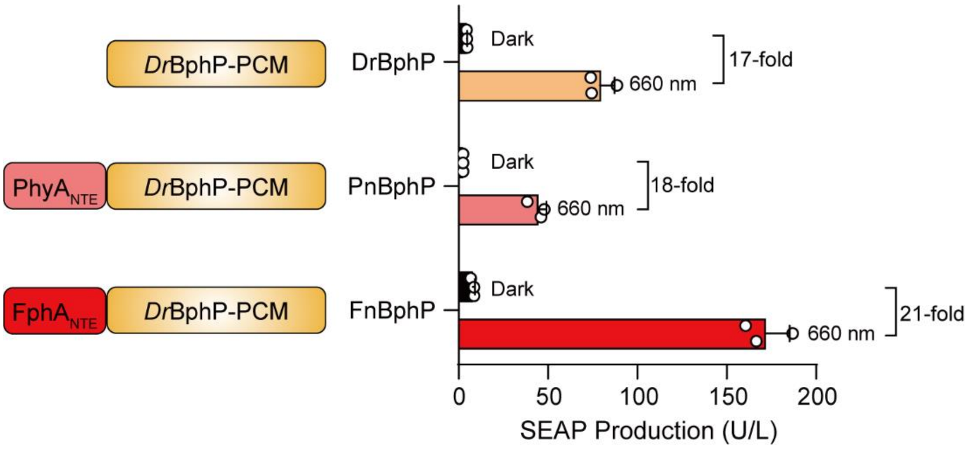


**Supplementary Fig. 1 Performance comparison between the different chimeric BphP constructs fused to the Gal4 domain.** HEK-293T cells (6×10$^4$) were co-transfected with a SEAP reporter expression vector (pDL6), the LDB3-VP64 expression vector (pQL207), and the different chimeric

BphP: Gal4-*Dr*BphP expression vector (pQL217), Gal4-PnBphP expression vector (pQL325), or Gal4-FnBphP expression vector (pQL326), and then illuminated with red light (RL, 660 nm, 2.0 mW/cm$^2$) for 24 hours; SEAP production in the culture supernatant was quantified 24 hours after illumination. Data are presented as means ± SD; *n* = 3 independent experiments. Detailed descriptions of the genetic constructs and transfection mixtures are provided in Supplementary Tables 1 and 5.

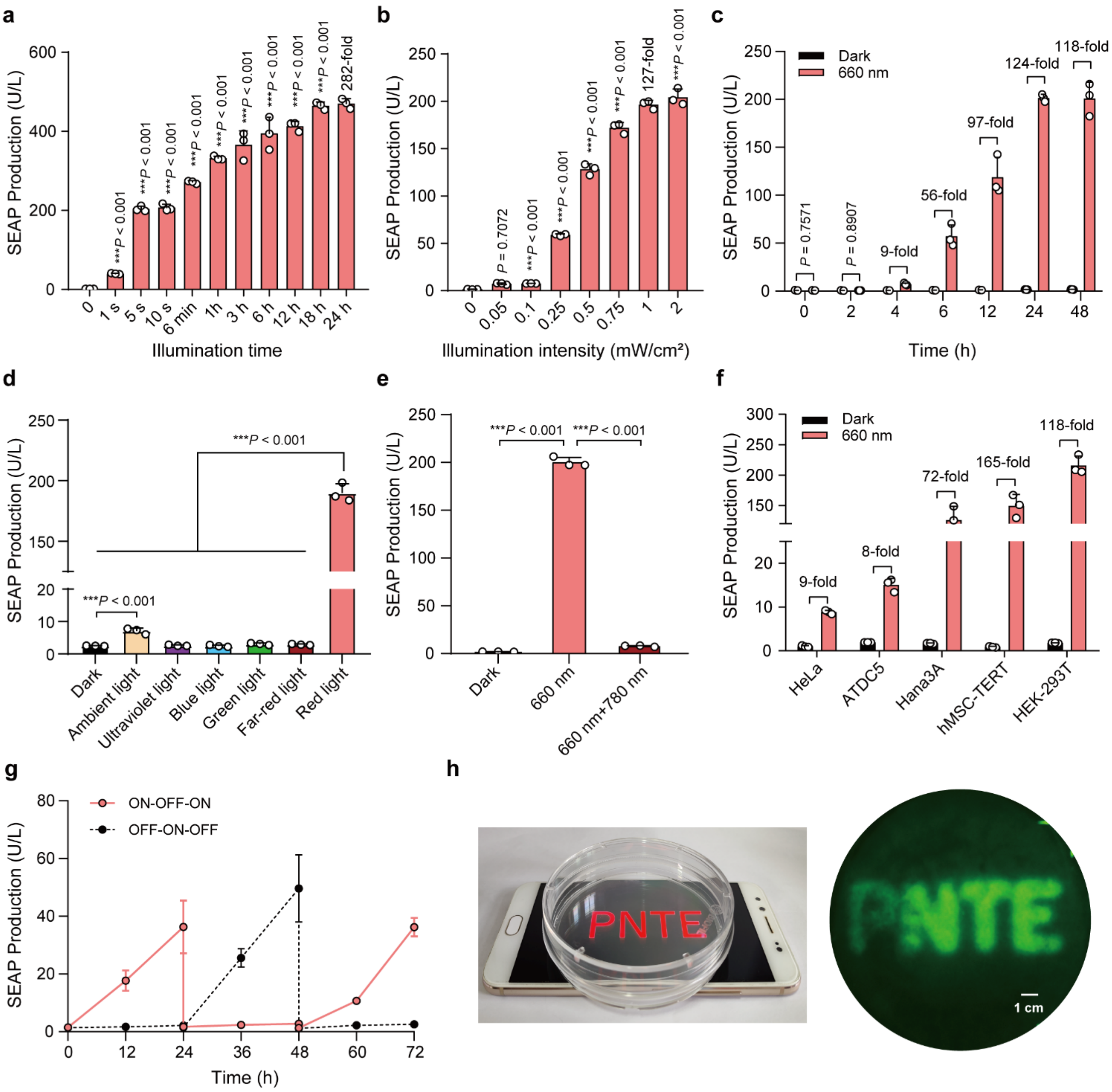


**Supplementary Fig. 2 Characterization of the performance of the Pn-REDLIP system in mammalian cells. a,** Assessment of illumination time-dependent Pn-REDLIP-mediated transgene expression kinetics. HEK-293T cells ($6\times10^4$) co-transfected with Gal4-PnBphP vector (pQL325), LDB3-p65-HSF1 expression vector (pNX12), and the SEAP reporter expression vector (pYZ430, $P_{RL}$-SEAP-pA) at a 2:2:1 (w/w/w) ratio were illuminated with RL (660 nm, 2.0 mW/cm$^2$) for the indicated time (0-24 hours). SEAP production was quantified 24 hours after initial illumination. **b,** Illumination intensity-dependent Pn-REDLIP-mediated transgene expression kinetics. HEK-293T cells ($6\times10^4$) transfected as described in a were illuminated with RL (660 nm) at the indicated light intensities (0 to 2.0 mW/cm$^2$) for 10 seconds. SEAP production was quantified 24 hours after illumination. **c,**

Quantification of Pn-REDLIP-mediated transgene expression kinetics. HEK-293T cells ($6\times10^4$) transfected as described in a were illuminated with RL (660 nm, 1.0 mW/cm$^2$) for 10 seconds. After initial illumination, SEAP production in the culture supernatant was profiled at the indicated time periods (0-48 hours). **d,** Chromatic specificity of the Pn-REDLIP system. HEK-293T cells ($6\times10^4$) transfected as described in a were illuminated with ultraviolet light (365 nm), blue light (465 nm), green light (530 nm), far-red light (780 nm), or red light (660 nm) at 1.0 mW/cm$^2$ or ambient light (750 Lux) for 10 seconds. SEAP production was quantified 24 hours after illumination. **e,** The switch ON/OFF performance of the Pn-REDLIP system. Three groups of HEK-293T cells were co-transfected as described in a. 24 hours after transfection, cells were illuminated with RL for 10 seconds and switched to the dark condition or 780 nm illumination at 2.0 mW/cm$^2$ for 2 minutes for the "660 nm+780 nm" group. As a control, cells were kept in the dark throughout the experiment. SEAP production was quantified at 24 hours after 660 or 780 nm illumination. **f,** Pn-REDLIP-mediated SEAP expression in the indicated mammalian cell lines. Five mammalian cell lines ($6\times10^4$) co-transfected as described in a were illuminated with RL (660 nm, 1.0 mW/cm$^2$) for 10 seconds. SEAP production was quantified 24 hours after illumination. **g,** Reversibility of Pn-REDLIP-mediated transgene expression. HEK-293T cells ($6\times10^4$) were transfected as described in (A) and were illuminated with RL (660 nm, 1.0 mW/cm$^2$) for 3 seconds (ON) or kept in the dark (OFF). SEAP production was quantified every 12 hours for 72 hours; the culture medium was renewed every 24 hours. **h,** Spatial control of Pn-REDLIP-mediated transgene expression. HEK-293T cells ($3\times10^6$) were seeded into a 10-cm tissue culture dish and transfected with the Pn-REDLIP system and an EGFP reporter pDQ63 ($P_{RL}$-EGFP-pA). The dish was placed onto a smartphone screen projecting a letter pattern and illuminated with RL for 15 minutes (display brightness set to 100%, 40 μW/cm$^2$) (schematic, left). The fluorescence micrographs assessing EGFP production were taken at 24 hours after illumination. Scale bar, 1 cm. Data in a-g are presented as means ± SD; $n$ = 3 independent experiments. $P$ values in (a, b, d, and e) were calculated by one-way ANOVA with multiple comparisons. $P$ values in c were calculated using a two-tailed unpaired $t$-test. Detailed descriptions of the genetic constructs and transfection mixtures are provided in Supplementary Tables 1 and 5.

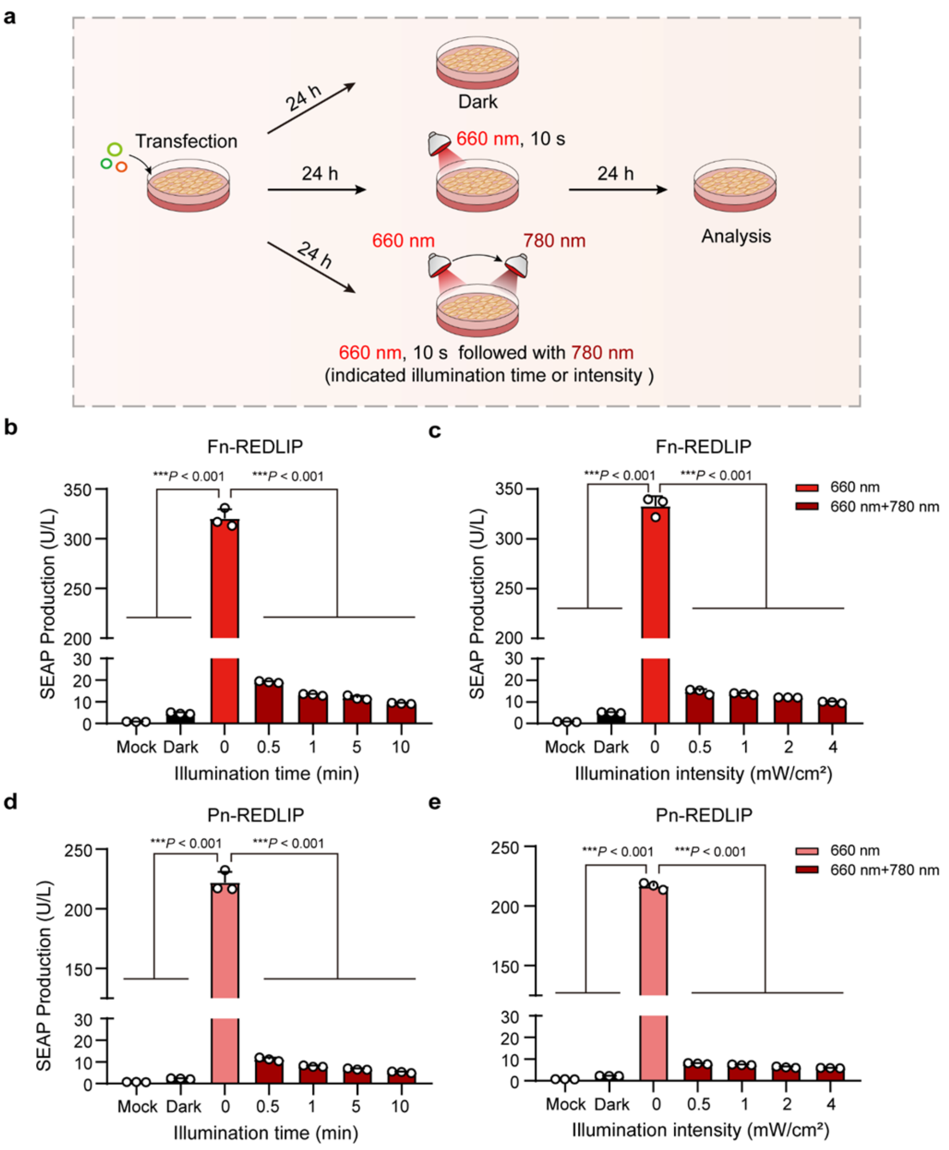


**Supplementary Fig. 3 Characterization of far-red light-dependent OFF kinetics of the Fn-REDLIP and Pn-REDLIP systems. a,** Schematic for the time schedule and experimental procedure of the activation/deactivation performance of the Fn-REDLIP and Pn-REDLIP systems. HEK-293T cells were co-transfected with the Fn-REDLIP or Pn-REDLIP systems and were illuminated with RL (660 nm, 2.0 mW/cm$^2$) for 10 seconds or maintained in the dark. Immediately, cells illuminated with RL were exposed to FRL (780 nm) for different time (0 to 10 minutes) or various light intensities (0 to 4.0 mW/cm$^2$). SEAP expression in the culture supernatant was quantified 24 hours after illumination. **b,** Illumination time-dependent Fn-REDLIP-mediated OFF kinetics of transgene expression. HEK-293T cells (6×10$^4$) were co-transfected with pQL326, pNX12, and pYZ430, and illuminated with RL (660 nm, 2.0 mW/cm$^2$) for 10 seconds or maintained in the dark. Immediately, cells illuminated with RL were exposed to FRL (780 nm, 1.0 mW/cm$^2$) for the indicated times. SEAP expression in the culture supernatant was quantified 24 hours after illumination. **c,** Illumination intensity-dependent Fn-REDLIP-mediated OFF kinetics of transgene expression. HEK-293T cells ($6 \times 10^4$) were co-transfected with pQL326, pNX12, and pYZ430, and illuminated with RL (660 nm, 2.0 mW/cm$^2$) for 10 seconds or maintained in the dark. Immediately, cells illuminated with RL were exposed to FRL (780 nm) of various light intensities (0 to 4.0 mW/cm$^2$) for 1 minute. SEAP expression in the culture supernatant was quantified 24 hours after illumination. **d,** Illumination time-dependent Pn-REDLIP-mediated OFF kinetics of transgene expression. HEK-293T cells ($6 \times 10^4$) were co-transfected with pQL325, pNX12, and pYZ430, and illuminated with RL (660 nm, 2.0 mW/cm$^2$) for 10 seconds or maintained in the dark. Immediately, cells illuminated with RL were exposed to FRL (780 nm, 1.0 mW/cm$^2$) for different time (0 to 10 minutes). SEAP expression in the culture supernatant was quantified 24 hours after illumination. **e,** Illumination intensity-dependent Pn-REDLIP-mediated OFF kinetics of transgene expression. HEK-293T cells ($6 \times 10^4$) were co-transfected with pQL325, pNX12, and pYZ430, and illuminated with RL (660 nm, 2.0 mW/cm$^2$) for 10 seconds or maintained in the dark. Immediately, cells illuminated with RL were exposed to FRL (780 nm) of various light intensities (0 to 4.0 mW/cm$^2$) for 1 minute. SEAP expression in the culture supernatant was scored 24 hours after illumination. Data are presented as means ± SD; $n = 3$ independent experiments. $P$ values were calculated by one-way ANOVA with multiple comparisons. Detailed descriptions of genetic components and transfection mixtures are provided in Supplementary Tables 1 and 5.

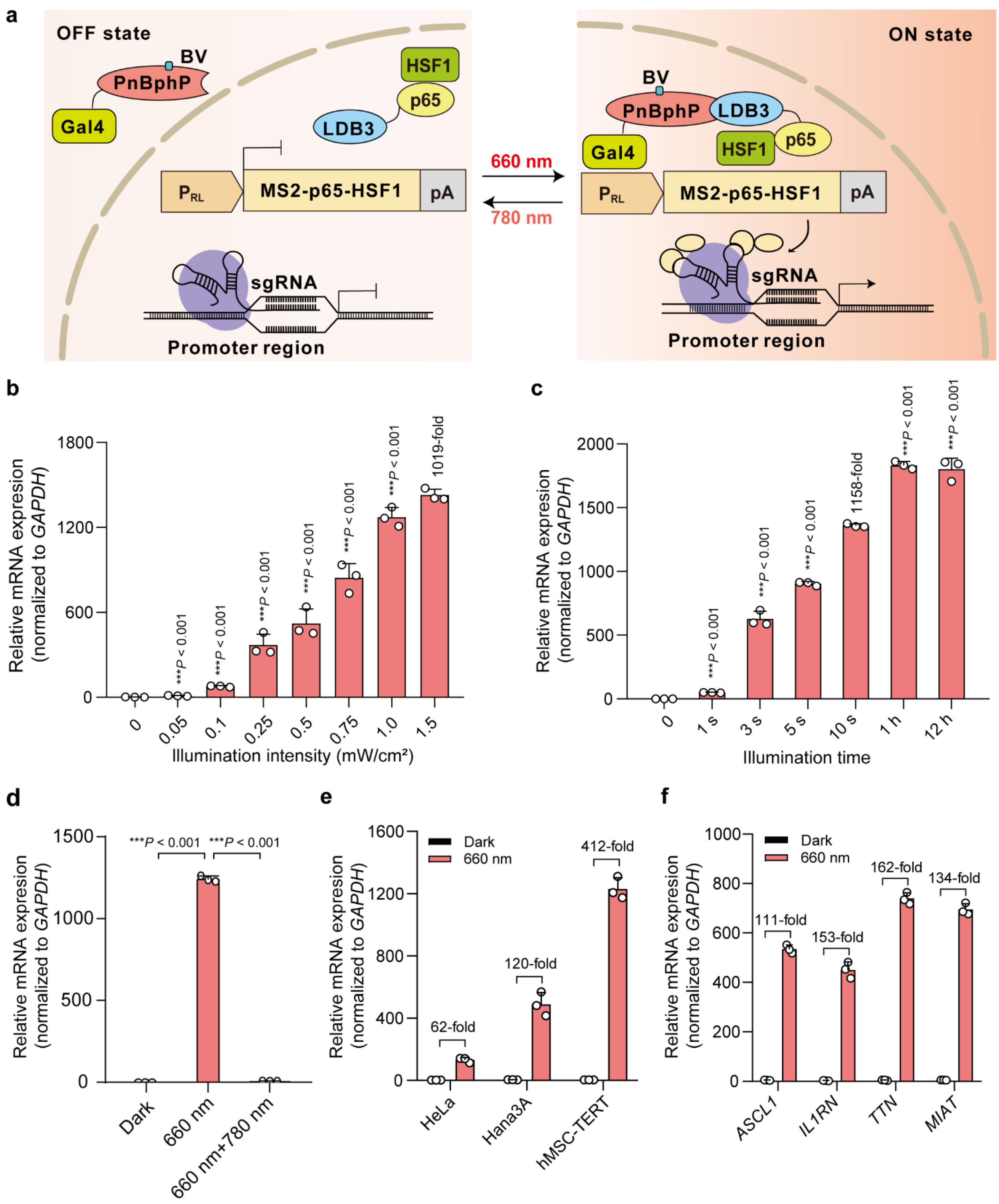


**Supplementary Fig. 4 Pn-REDLIP-controlled CRISPR-dCas9 system (Pn-REDLIP$_{cas}$) for endogenous gene activation *in vitro*. a,** Schematic design of the Pn-REDLIP$_{cas}$ system for transcriptional activation. The RL chimeric promoter (P$_{RL}$) drove the expression of the *trans*-activator MS2-p65-HSF1, which can be recruited by the MS2 box of the sgRNA-dCas9 complex to activate target gene transcription under RL (660 nm) illumination. FRL (780 nm) illumination can switch the Pn-REDLIP$_{cas}$ system back to an inactive state. **b,** Illumination intensity-dependent Pn-REDLIP$_{cas}$ system-mediated activation of endogenous gene transcription. HEK-293T ($6\times10^4$) cells were co-transfected with pQL325, pNX12, a P$_{RL}$-driven MS2-p65-HSF1 expression vector (pDQ100), dCas9 expression vector (pSZ69), and two sgRNAs targeting the RHOXF2 locus (pSZ105, P$_{U6}$-sgRNA1$_{RHOXF2}$-pA; pSZ106, P$_{U6}$-sgRNA2$_{RHOXF2}$-pA) at a 15:15:1:10:5:5 (w/w/w/w/w/w) ratio, followed by illumination with RL (660 nm) at the indicated intensities (0 to 1.5 mW/cm$^2$) for 10 seconds. **c,** Illumination time-dependent Pn-REDLIP$_{cas}$ system-mediated activation of endogenous gene transcription. HEK-293T cells ($6\times10^4$) were co-transfected as described in b and illuminated with RL (660 nm, 1.0 mW/cm$^2$) for different time periods (0-12 hours). **d,** The switch ON/OFF performance of the Pn-REDLIP$_{cas}$ system. Three groups of HEK-293T cells were transfected as described in b. 24 hours after transfection, cells were illuminated with RL for 10 seconds and transitioned to the dark condition or FRL illumination at 2.0 mW/cm$^2$ for 2 minutes. As a control, cells were kept in the dark throughout the experiment. The levels of endogenous *RHOXF2* were quantified by qPCR at 24 hours after 660 nm or 780 nm illumination. **e,** The Pn-REDLIP$_{cas}$-mediated endogenous gene activation in the indicated mammalian cell lines. Three mammalian cell lines ($6\times10^4$) were co-transfected as described in b and illuminated with RL (660 nm, 1.0 mW/cm$^2$) for 10 seconds. **f,** The Pn-REDLIP$_{cas}$-mediated activation of different endogenous genes. HEK-293T cells ($6\times10^4$) were co-transfected with Pn-REDLIP$_{cas}$ together with two sgRNAs targeting the *ASCL1* locus, the *TTN* locus, the *IL1RN* locus, or the *MIAT* locus. Twenty-four hours after transfection, cells were exposed to RL (660 nm, 1.0 mW/cm$^2$) for 10 seconds. Data in b-f are presented as relative mRNA expression levels quantified by qPCR 24 hours after illumination. All data are represented as means ± SD; $n$ = 3 independent experiments. $P$ values were calculated by one-way ANOVA with multiple comparisons. Detailed descriptions of genetic components and transfection mixtures are provided in Supplementary Tables 1 and 5.

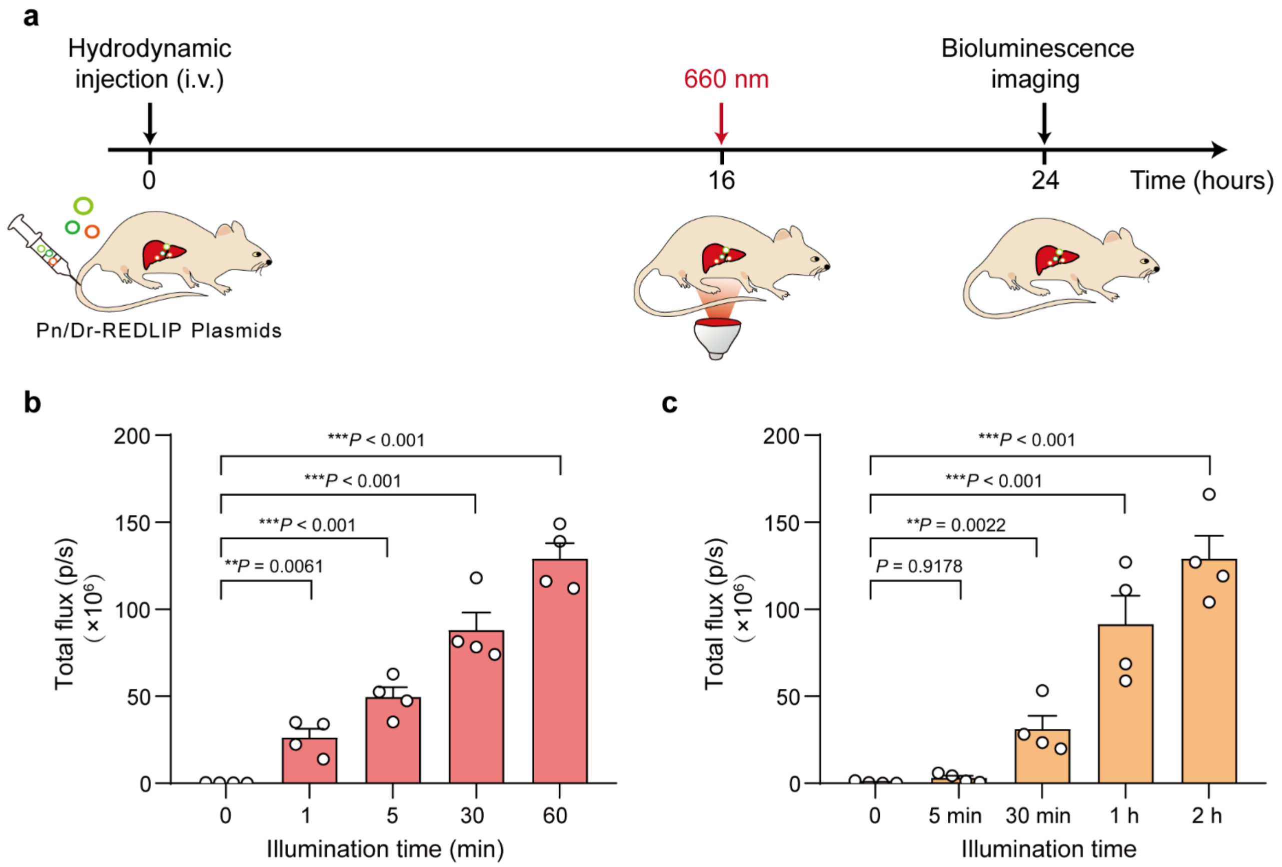


**Supplementary Fig. 5 Pn-REDLIP and Dr-REDLIP mediated transgene expression in mice. a,** Schematic representation of the experimental procedure and time schedule for Pn-REDLIP and Dr-REDLIP mediated transgene expression in mice. **b,** Exposure time-dependent Pn-REDLIP-mediated transgene expression kinetics in mice. Mice hydrodynamically injected with Pn-REDLIP-encoding plasmids were illuminated with RL (660 nm, 20 mW/cm$^2$) for different time periods. The bioluminescence signal was quantified 8 hours after illumination using an *in vivo* imaging system. **c,** Exposure time-dependent Dr-REDLIP-mediated transgene expression kinetics in mice. Mice hydrodynamically injected with Dr-REDLIP-encoding plasmids were illuminated with RL (660 nm, 20 mW/cm$^2$) for different time periods. The bioluminescence signal was quantified 8 hours after illumination using an *in vivo* imaging system. Data are presented as means ± SEM ($n$ = 4 mice). $P$ values were calculated by one-way ANOVA with multiple comparisons. Detailed bioluminescence images of the mice are provided in Supplementary Fig. 6b, c.

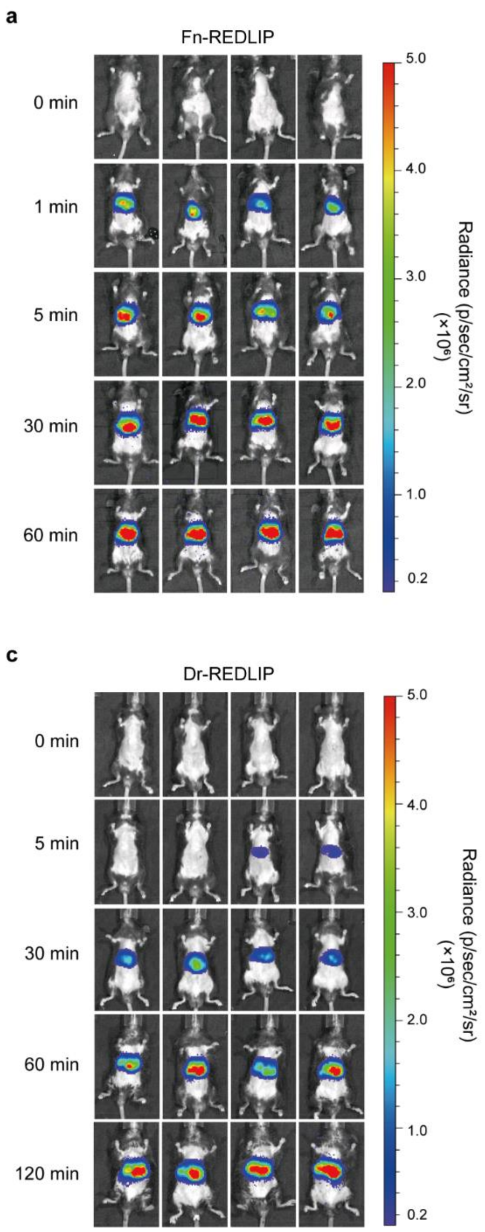


**Supplementary Fig. 6 Illumination of three types of time-dependent REDLIP systems-mediated transgene expression kinetics in mice (related to Fig. 4c and Supplementary Fig. 6b, c).** *In vivo* bioluminescence images of mice illuminated with RL for different time periods. Mice hydrodynamically injected with three types of REDLIP system-encoded plasmids were illuminated with RL (660 nm, 20 mW/cm$^2$) for different time periods, and bioluminescence was quantified at 8 hours after illumination. (a) Fn-REDLIP; (b) Pn-REDLIP; (c) Dr-REDLIP). $n$ = 4 mice.

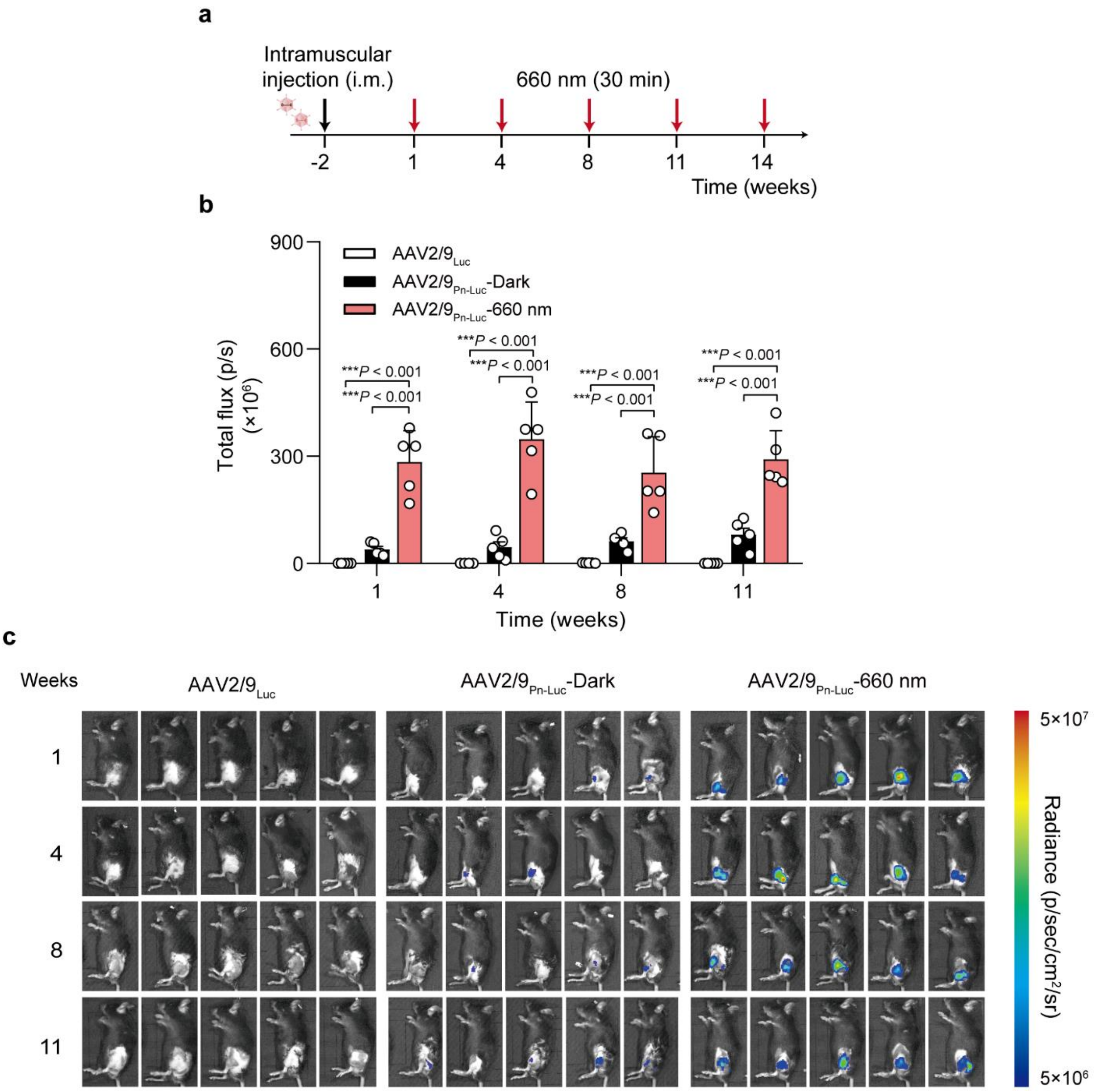


**Supplementary Fig. 7 Pn-REDLIP-mediated transgene expression in mice. a,** Schematic representation of the experimental procedure and time schedule for the AAV-2/9 delivery of Pn-REDLIP-mediated transgene expression in the mouse left gastrocnemius muscle. Mice were intramuscularly injected with a mixture of AAV2/9 vectors encoding the Pn-REDLIP system [(pNX257 (ITR-$P_{EMS}$-Gal4-PnBphP-pA-ITR)，pNX137 (ITR-$P_{EMS}$-LDB3-p65-HSF1-pA-ITR)] and the luciferase reporter pQL271 (ITR-$P_{RL}$-Luciferase-pA-ITR) or only the luciferase reporter pQL271 at an AAV titer of $2\times10^{11}$ vg. The control group received an intramuscular injection (left gastrocnemius muscle) of the RL-responsive luciferase reporter ($AAV2/9_{Luc}$, $2\times10^{11}$ vg). After 2 weeks, the injected mice were illuminated at an intensity of 20 mW/cm$^2$ for 30 minutes once every three or four weeks. **b,**

Bioluminescence was quantified 8 hours after illumination. **c,** Bioluminescence measurements of Pn-REDLIP-mediated luciferase expression in AAV-transduced mice under RL illumination. Data in b are presented as means ± SEM (*n* = 5 mice). *P* values in b were calculated by one-way ANOVA with multiple comparisons.

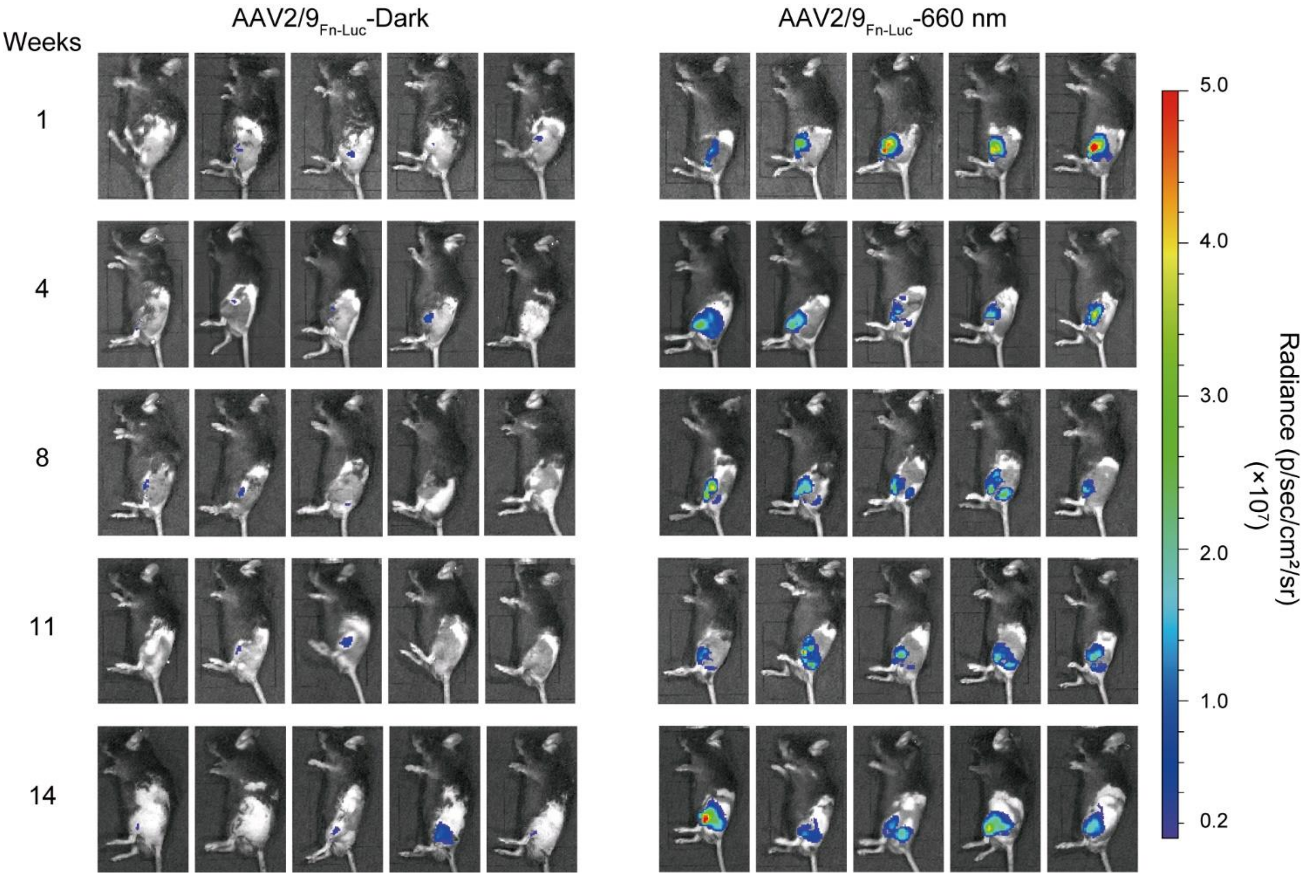


**Supplementary Fig. 8 Long-term study of AAV-delivered Fn-REDLIP-mediated transgene expression in mice (related to Fig. 4e).** A mixture of AAV-2/9 vectors encoding Fn-REDLIP systems were intramuscularly injected into the left gastrocnemius muscle of mice. Two weeks after the AAV transduction, mice were illuminated with RL (20 mW/cm$^2$) at an intensity of 20 mW/cm$^2$ for 30 minutes once every three or four weeks. Bioluminescence was quantified 8 hours after illumination. $n$ = 5 mice.

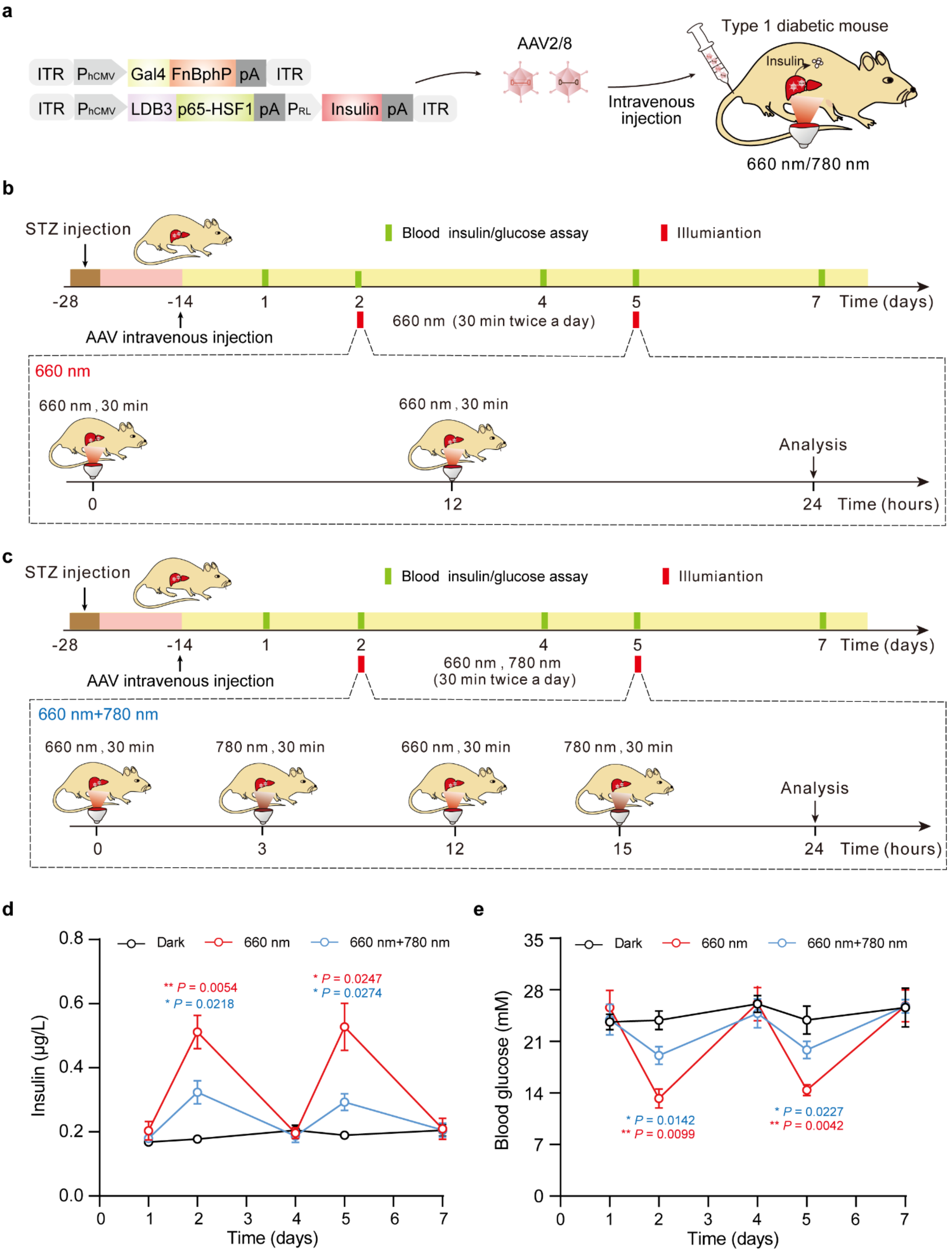


**Supplementary Fig. 9 AAV-delivered Fn-REDLIP system controlling hepatic insulin expression in T1D mice with on/off capability. a,** Schematic for the AAV-2/8 vectors for the Fn-REDLIP system iteration containing pNX177 (ITR-P$_{hCMV}$-Gal4-FnBphP-pA-ITR) and the concatenated vector pQL318 (ITR-$P_{hCMV}$-LDB3-p65-HSF1-pA::$P_{RL}$-insulin-pA-ITR). **b, c,** Schematic illustration for the experimental procedure and time schedule for AAV-delivered Fn-REDLIP-mediated insulin expression in T1D model mouse livers. T1D mice were intravenously injected with a mixture of AAV-2/8 vectors (targeting the livers) encoding the Fn-REDLIP system. After 2 weeks, the injected T1D mice were randomly divided into three groups: i) Dark group: T1D mice kept in the dark; ii) 660 nm group: T1D mice exposed to RL (660 nm, 20 mW/cm$^2$) for 30 minutes twice a day; iii) 660 nm+780 nm group: T1D mice exposed to RL (660 nm, 20 mW/cm$^2$) for 30 minutes twice a day (ON), followed by exposure to FRL (780 nm, 20 mW/cm$^2$) for 30 minutes twice a day (OFF) after a 3-hour interval. **d,** Blood insulin was profiled at the indicated time points using a mouse insulin ELISA kit. **e,** Blood glucose was profiled at the indicated time points using a blood glucose meter. Data in d, e are presented as means ± SEM ($n$ = 5 mice). $P$ values in d, e were calculated by one-way ANOVA with multiple comparisons. $P$ values in d, e were calculated by comparing the 660 nm group or the 660 nm+780 nm group with the Dark group.

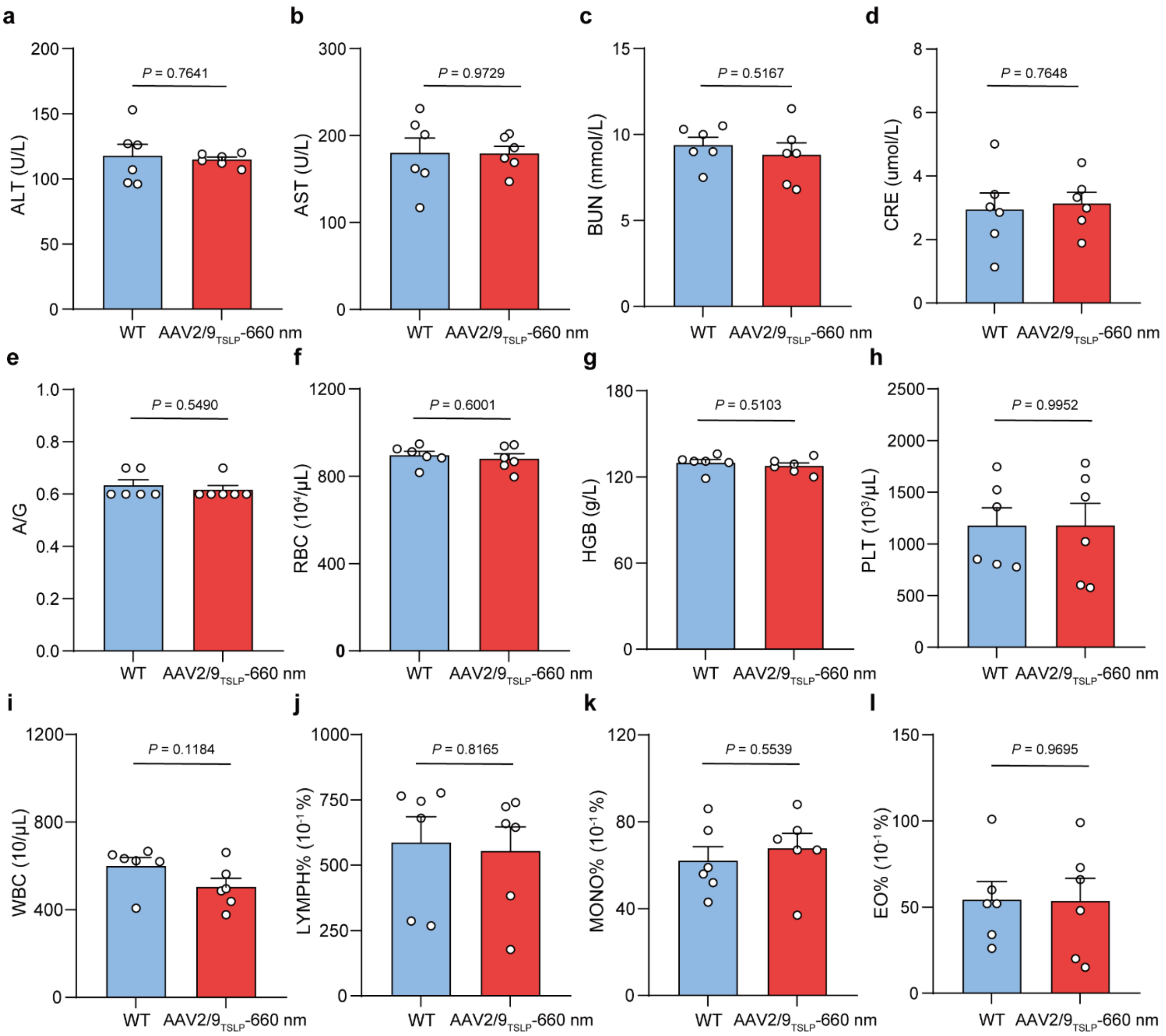


**Supplementary Fig. 10 Blood biochemistry and hematology assay of HFD mice treated with AAV-delivered Fn-REDLIP and healthy wild type (WT) C57BL/6 mice for two months.** HFD mice were intramuscularly injected in the left gastrocnemius muscle with a mixture of AAV-2/9 vectors encoding Fn-REDLIP. After 2 weeks, the injected HFD mice were illuminated with RL at an intensity of 20 mW/cm$^2$ for 30 minutes every three days for 6 weeks (AAV2/9$_{TSLP}$-660 nm). The examined control was non-model wild-type mice. Eight weeks after AAV transduction, mouse blood was collected to quantify the (**a**) glutamic-pyruvic transaminase (ALT), (**b**) aspartate aminotransferase (AST), (**c**) blood urea nitrogen (BUN), (**d**) creatinine (CRE), (**e**) albumin/globulin ratio (A/G), (**f**) counts for red blood cells (RBC), (**g**) hemoglobin (HGB), (**h**) platelet (PLT), (**i**) total white blood cells (WBC), (**j**) percentage of blood lymphocytes (LYMPH), (**k**) blood monocytes (MONO), and (**l**) blood eosinophil (EO). All data are expressed as means ± SEM ($n$ = 6 mice). $P$ values were calculated using a two-tailed unpaired $t$-test.

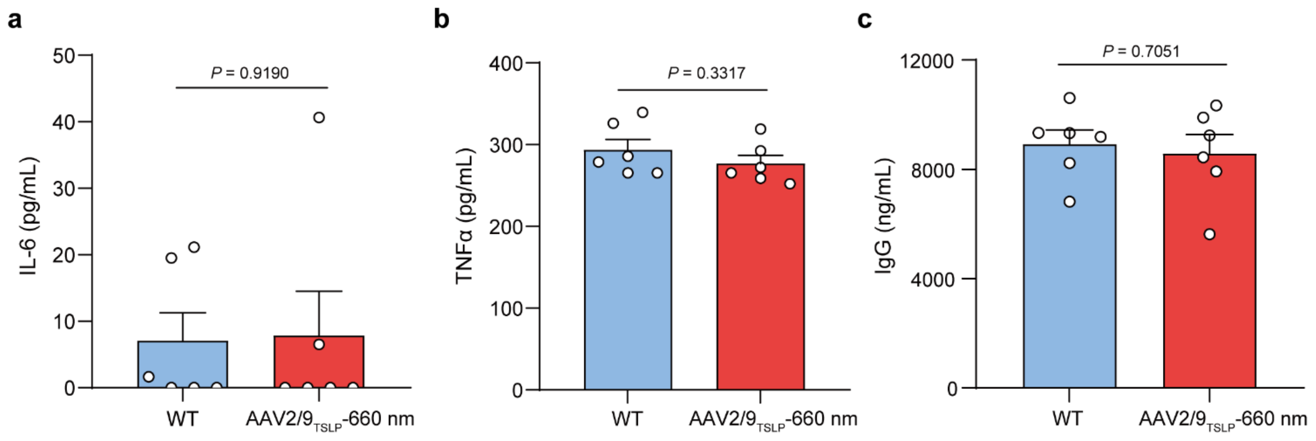


**Supplementary Fig. 11 Serum inflammatory cytokines and immunoglobulin in HFD mice treated with AAV-delivered Fn-REDLIP and wild-type C57BL/6 mice for two months.** HFD mice were intramuscularly injected in the left gastrocnemius muscle with a mixture of AAV-2/9 vectors encoding Fn-REDLIP. Two weeks after AAV transduction, the injected HFD mice were illuminated with RL at an intensity of 20 mW/cm$^2$ for 30 minutes every three days for 6 weeks (AAV2/9$_{TSLP}$-660 nm). The examined control was non-model wild-type mice. Eight weeks after the AAV transduction, mouse blood was collected to quantify (**a**) serum IL-6, (**b**) serum TNF-α, and (**c**) serum IgG production by ELISA. All data are expressed as means ± SEM ($n$ = 6 mice). $P$ values were calculated using a two-tailed unpaired $t$-test.

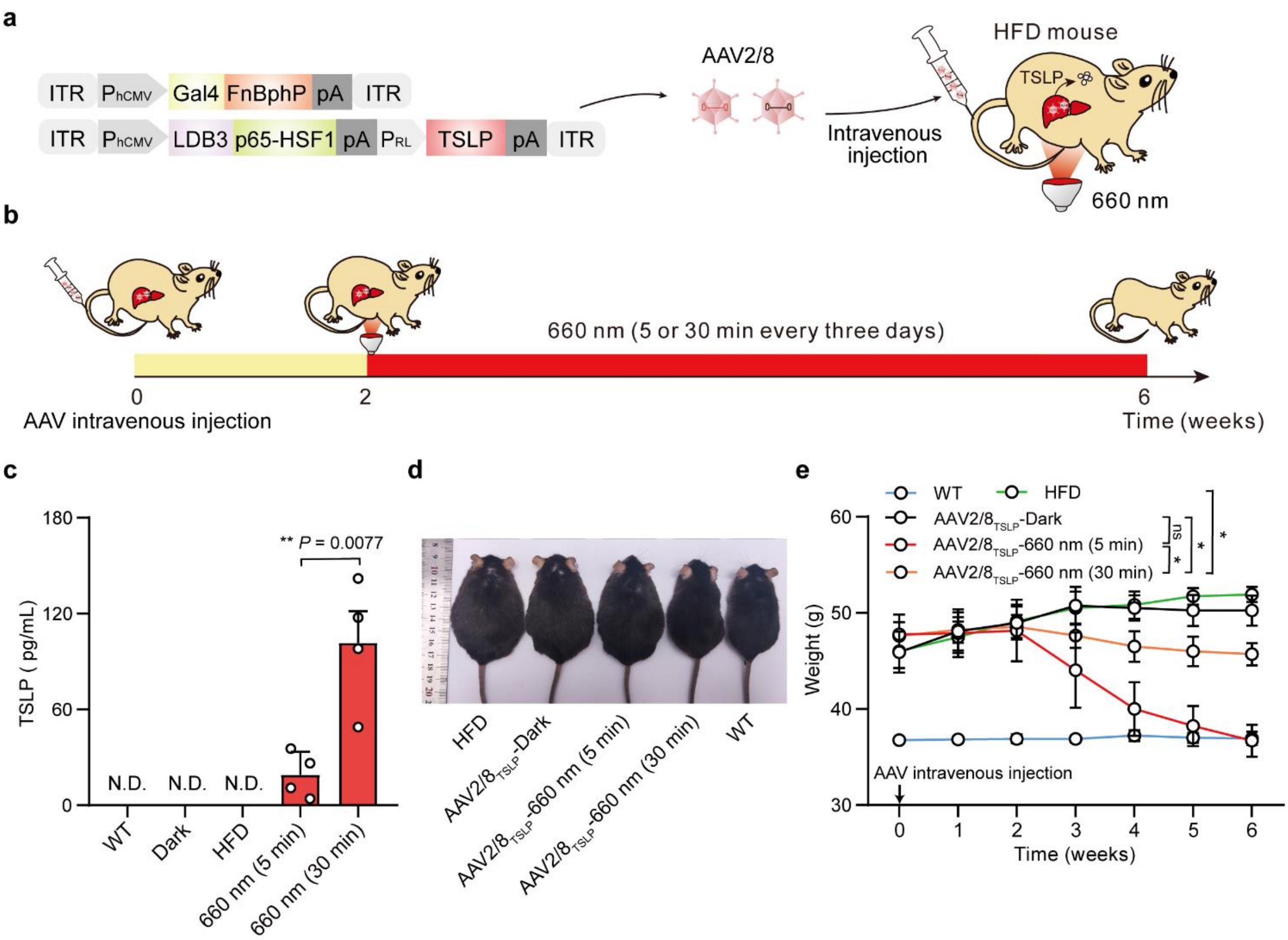


**Supplementary Fig. 12 AAV-delivered Fn-REDLIP system for hepatic TSLP expression to regulate body weight in HFD mice. a,** Schematic for the genetic configuration of the AAV-2/8 vectors for the Fn-REDLIP system used for the HFD obesity model mice experiment. **b,** Schematic for the experimental procedure and time schedule for AAV-delivered Fn-REDLIP-mediated TSLP transgene expression in HFD obesity model mouse livers. HFD mice were intravenously injected with a mixture of AAV-2/8 vectors (targeting the livers) encoding the Fn-REDLIP system iteration containing pNX177 (ITR-$P_{hCMV}$-Gal4-FnBphP-pA-ITR) and the concatenated vector pNX166 (ITR-$P_{hCMV}$-LDB3-p65-HSF1-pA::$P_{RL}$-TSLP-pA-ITR) at an AAV titer of $2\times10^{11}$ vg. After 2 weeks, the injected HFD mice were illuminated with RL (20 mW/cm$^2$) for 5 or 30 minutes every three days for 4 weeks (AAV2/8$_{TSLP}$-660 nm). The examined controls included non-model wild-type control mice (WT group), untreated HFD control mice (HFD group), and AAV-injected HFD mice kept in the dark (AAV2/8$_{TSLP}$-Dark). **c,** The levels of TSLP in serum were quantified using a mouse TSLP ELISA kit. N.D., Not detected. **d,** Representative images of the AAV2/8$_{TSLP}$-660 nm group at 6 weeks after AAV injection. **e,** Analysis of body weight of HFD mice injected with the AAV2/8-delivered Fn-REDLIP system. ns, not significant, *$P$ = 0.0370, *$P$ = 0.0242, **$P$ = 0.0134. Data in c, e are presented as means ± SEM ($n$ = 4 mice). $P$ values in c were calculated using a two-tailed unpaired $t$-test. $P$ values in e were calculated by one-way ANOVA with multiple comparisons.

**Supplementary Table 1. Plasmids designed and used in this study**

**Abbreviations**: **AAV**, adeno-associated virus; **Aff6-V18FΔN**, a photo-state-specific binder; ***ASCL1*,** human achaete-scute homolog 1 gene; ***Ascl1*,** mouse achaete-scute homolog 1 gene; **BAT,** brown adipose tissue; **BldD**, *Streptomyces coelicolor* transcription factor regulating aerial hyphae formation; **BphP1**, *Rhodopseudomonas palustris* bacteriophytochrome; **eWAT,** epididymal white adipose tissue; **DrBphP**, a photosensory protein; **EGFP**, enhanced green fluorescent protein; **Fd**, ferredoxin; **FHY1**, far red elongated hypocotyl 1; **FnBphP,** chimeric photosensory protein fusing the N-terminal extension (NTE) of FphA to N-terminus of *Dr*BphP-PCM; **FNR**, ferredoxin-NADP$^+$ reductase; **Gal4**, galactose-responsive transcription factor 4; **HO1**, heme oxygenase; ***IL1RN***, interleukin 1 receptor antagonist gene; **ITR**, inverted terminal repeat; **iWAT,** inguinal white adipose tissue; **LDB3**, one light-form specific nanobody; **Luciferase**, bioluminescence enzymes; ***MIAT***, myocardial infarction associated transcript; **MS2**, an RNA aptamer; **MTAD**, 5-methylthioadenosine/S-adenosylhomocysteine deaminase; **NLS**, mammalian nuclear localization signal; **P2A**, picornavirus-derived self-cleaving peptide engineered for bicistronic gene expression in mammalian cells; **p65**, 65kDa transactivator subunit of NF-kB; **p65-HSF1**, fused transactivator of P65 and heat shock factor 1(HSF1); **pA**, polyadenylation signal; **PcyA**, ferredoxin oxidoreductase; **$P_{EMS}$**, enhanced skeletal muscle cell specific promoter; **$P_{hCMV}$**, human cytomegalovirus immediate early promoter; **$P_{hCMVmin}$**, minimal version of $P_{hCMV}$; **ΔPhyA**, one type of truncated PhyA(1-617aa); **PhyB**, phytochrome B; **PIF3**, phytochrome interacting factor 3; **PnBphP,** chimeric photosensory protein fusing the N-terminal extension (NTE) of PhyA to N-terminus of *Dr*BphP-PCM; **PpsR2**, the natural cognate repressor of BphP1; ***RHOXF***, rhox homeobox family member 1 gene; **SEAP**, secreted alkaline phosphatase; **sgRNA**, single guide RNA; **TSLP,** Thymic stromal lymphopoietin induces; ***TTN***, titin gene; **UAS**, Gal4-specific binding sequence; **VP16**, a *Herpes simplex* virus (HSV)-derived transcriptional activator protein; **VP64**, a tetrameric repeat of the minimal activation domain derived from the *Herpes simplex* virus protein VP16; **VPR**, a tripartite activator VP64-p65-Rta fusion protein; **whiG**, BldD-specific binding sequence; **YhjH,** a bacterial c-di-GMP phosphodiesterase.

**Supplementary Table 2. Target sequences of sgRNAs used in this study**

| Gene name | sgRNA name | Guide sequence of sgRNA (5'-3') |
|---|---|---|
| *RHOXF2* (human) | sgRNA1$_{RHOXF2}$ | ACGCGTGCTCTCCCTCATC |
| | SgRNA2$_{RHOXF2}$ | CTGTGGGTTGGGCCTGCTG |
| *ASCL1* (human) | sgRNA1$_{ASCL1}$ | GGCTGGGTGTCCCATTGAAA |
| | SgRNA2$_{ASCL1}$ | ATGGAGAGTTTGCAAGGAGC |
| *IL1RN* | sgRNA1$_{IL1RN}$ | TGTACTCTCTGAGGTGCTC |

| | | |
|---|---|---|
| (human) | SgRNA2$_{IL1RN}$ | GAGTCACCCTCCTGGAAAC |
| *TTN* (human) | sgRNA1$_{TTN}$ | CCTTGGTGAAGTCTCCTTTG |
| | SgRNA2$_{TTN}$ | ATGTTAAAATCCGAAAATGC |
| *MIAT* (human) | sgRNA1$_{MIAT}$ | GCGCCCATGAAATTTTAATG |
| | SgRNA2$_{MIAT}$ | GCTTCTGCGCCCCTGGTCCG |
| *Ascl1* (mouse) | sgRNA1$_{Ascl1}$ | GCAGCCGCTCGCTGCAGCAG |
| | SgRNA2$_{Ascl1}$ | AGCTGAGGAGGTGGGGGAAG |

**Supplementary Table 3. Oligonucleotide sequences used for qPCR analysis**

| Gene name | Primer name | Primer sequence (5'-3') |
|---|---|---|
| *GAPDH* (human) | Forward | CGAGATCCCTCCAAAATCAA |
| | Reverse | ATCCACAGTCTTCTGGGTGG |
| *RHOXF2* (human) | Forward | AGTGTAGCCAGTATATGACCAGC |
| | Reverse | TGACCTCTTCAGTAAGCGACAG |
| *ASCL1* (human) | Forward | CGCGGCCAACAAGAAGATG |
| | Reverse | CGACGAGTAGGATGAGACCG |
| *IL1RN* (human) | Forward | CATTGAGCCTCATGCTCTGTT |
| | Reverse | CGCTGTCTGAGCGGATGAA |
| *TTN* (human) | Forward | CCCCATCGCCCATAAGACAC |
| | Reverse | CCACGTAGCCCTCTTGCTTC |
| *MIAT* (human) | Forward | TGGCTGGGGTTTGAACCTTT |
| | Reverse | AGGAAGCTGTTCCAGACTGC |
| *Gapdh* (mouse) | Forward | ATGACATCAAGAAGGTGGTG |
| | Reverse | CATACCAGGAAATGAGCTTG |
| *Ascl1* (mouse) | Forward | GGAACAAGAGCTGCTGGACT |
| | Reverse | GTTTTTCTGCCTCCCCATTT |

**Supplementary Table 4. Expression vectors and transfection mixtures used in Main Figures.**
